# GPI anchor remodeling by the plant PGAP1 ortholog HLD1 is essential for *Papaver* self-incompatibility

**DOI:** 10.1101/2021.05.20.444919

**Authors:** Zongcheng Lin, Fei Xie, Marina Triviño, Tao Zhao, Frederik Coppens, Lieven Sterck, Maurice Bosch, Vernonica E. Franklin-Tong, Moritz K. Nowack

**Author notes:** These authors contributed equally (Zongcheng Lin, Fei Xie, Marina Triviño). Corresponding authors Zongcheng Lin, Maurice Bosch, Vernonica E. Franklin-Tong, Moritz K. Nowack. Marina Triviño.

## Abstract

In eukaryotes, glycosylphosphatidylinositol anchored proteins (GPI-APs) are tethered to the outer leaflet of the plasma membrane where they function as key regulators of a plethora of biological processes. Self-incompatibility (SI) plays a pivotal role regulating fertilization in higher plants through recognition and rejection of ‘self’ pollen. Here we used *Arabidopsis thaliana* lines engineered to be self-incompatible by expression of *Papaver* SI determinants for an SI suppressor screen. We identify *HLD1*, an ortholog of human GPI-inositol deacylase *PGAP1*, whose mutation completely abolishes the SI response. We show that HLD1 functions as a GPI-inositol deacylase and that this GPI-remodeling activity is essential for SI. Using GFP-SKU5 as a representative GPI-AP, we show that *HLD1* mutation does not affect GPI-AP production and targeting, but alters the configuration of mature GPI-APs. This prevents GPI-AP release from the plasma membrane, suggesting that this process plays a critical role in the regulation of SI. Our data not only identify GPI anchoring as a new pathway of SI providing new directions to investigate SI mechanisms, but identifies for the first time a function for GPI-AP remodeling by inositol deacylation in plants.

**One sentence summary:** The *Papaver* self-incompatibility response requires GPI-anchor modification by HLD1, an ortholog of the mammalian PGAP1.

## Introduction

Glycosylphosphatidylinositol (GPI) anchoring is a highly conserved post-translational modification in eukaryotes and plays crucial roles in the targeting and tethering of GPI-anchored proteins (GPI-APs) to the outer leaflet plasma membrane^1,2^. Over 150 GPI-APs are predicted in humans and more than 250 in Arabidopsis (*Arabidopsis thaliana*)^3,4^. GPI-APs are functionally diverse; they are key regulators of cell signaling, growth, morphogenesis, reproduction and pathogenesis in yeast, mammals, and plants^5-7^. GPI-AP biosynthesis, remodeling, and export are well-characterized in yeast and animal cells^8,9^. The GPI anchor is synthesized and transferred to target proteins in the endoplasmic reticulum (ER)^1^. Knockouts of *Phosphatidylinositol glycan anchor biosynthesis* (*PIG*) genes in mouse and Arabidopsis show that GPI-anchor synthesis is essential for early embryo and tissue development^6,10,11^. Although comparatively little is known about this process in plants, studies have shown that several proteins involved in GPI anchor biosynthesis are required for plant viability and male fertility ^6^. Once the GPI anchor is attached to a target protein, several remodeling steps take place. In animals, the first step involves the removal of an acyl chain on the GPI inositol, by <u>p</u>ost-<u>G</u>PI <u>a</u>ttachment to <u>p</u>roteins <u>1</u> (PGAP1), a GPI-inositol deacylase^12^. This step is important for the efficient transport of GPI-APs to the outer leaflet of the plasma membrane ^5,8^, where they are either retained or released by cleavage^13^. However, although putative homologs of *PGAP1* have been identified in plant genomes^6,14^, there are no functional data about this GPI remodeling and maturation step in plants.

Self-incompatibility (SI) regulates the rejection of ‘self’ pollen that is deposited on the stigma of the same species, preventing inbreeding and promoting genetic diversity. Generally, SI is genetically controlled by a polymorphic multi-allelic *S*-locus, with each *S*-haplotype encoding a pair of *S*-determinants^15^. Stigmas discriminate between “self” and “non-self” pollen using allele-specific interactions between these two *S*-determinants. In field poppy (*Papaver rhoeas*), the female *S*-determinant PrsS is specifically expressed in the stigma^16^, whereas the male *S*-determinant PrpS is specifically expressed in pollen^17^. Cognate PrpS-PrsS interaction triggers a well characterized signaling network resulting in pollen tube growth arrest and programmed cell death (PCD) in ‘self’-pollen^18^. Recently, we showed that Arabidopsis cells (both vegetative and reproductive cells) expressing *PrpS* undergo an SI-PCD response comparable to *Papaver* pollen when treated with cognate recombinant PrsS protein *in vitro*^19,20^. Furthermore, cognate *PrpS* and *PrsS* co-expressed in the self-compatible model plant Arabidopsis trigger a robust *Papaver*-like SI response. This SI process rendered Arabidopsis lines effectively self-incompatible demonstrating that *Papaver S*-determinants are sufficient to confer functional SI in Arabidopsis^19,20^. These self-incompatible Arabidopsis lines (named *At-*SI hereafter) provide a facile macroscopic readout of SI-PCD, as self-pollinated *At*-SI lines do not set seed and silique development does not occur.

Here we used the *At-*SI lines as the basis for a forward genetic mutant screen to identify new genes involved in controlling *Papaver* SI. We identified 12 mutant alleles of *highlander1* (*hld1*) with suppressed SI resulting in restoration of self-compatibility. We demonstrated that HLD1 is a functional ortholog of the mammalian *PGAP1* gene involved in the deacylation of GPI-APs. The *HLD1* mutation results in “three-footed” GPI-APs but does not affect the expression of GPI-APs at the plasma membrane. This study reveals a crucial role of GPI-APs and GPI-remodeling of GPI-APs in the *Papaver* SI response and provides the first functional analysis of a GPI-remodeling protein in plants.

## Results

### Identification of SI-defective *highlander* mutants

To discover novel players of the *Papaver* SI process, we performed an ethyl methanesulfonate-(EMS-) based forward genetic screen on roughly 50,000 mutated *At*-SI M1 individuals for suppressors of SI (**Fig S1, Materials & Methods**). We identified twelve independent self-compatible SI-repressor mutants with high levels of self-seed set and silique development after excluding those mutants with mutations in the transgene or changes in the PrpS_1_-GFP expression (**Fig 1A; Fig S2)**. We named these mutants *highlander* (*hld*), after the immortal warriors in the 1980s cult film. To establish whether *hld* mutants disrupted SI signaling on the male (pollen) or female (stigma) side, we performed reciprocal crosses. Pollinating *hld* pollen onto *At-*SI stigmas resulted in significantly longer siliques (**Fig 1B**, p<0.0001) and higher seed set (**Fig S2C**, p<0.0001) than in the *At-*SI parent line, whereas the reciprocal cross did not (**Fig 1B**, p=0.1458, N.S.; **Fig S2C**, p=0.0909, N.S.). Thus, the *hld* mutations affect SI in the pollen but not in the stigma. The identification of *hld* mutants from the M1 generation after mutagenesis suggested the gametophytic nature of the *hld* mutations. This was further confirmed by the observation that after the first backcross (BC) to the *At*-SI parent line, all the heterozygous *hld* BC1 generation plants were self-compatible (**Table S1**, n= 167), demonstrating that the 50% mutant pollen grains produced by heterozygous plants is sufficient to restore seed set, showing that the *hlds* are gametophytic (pollen) mutants.

**Fig. 1.**
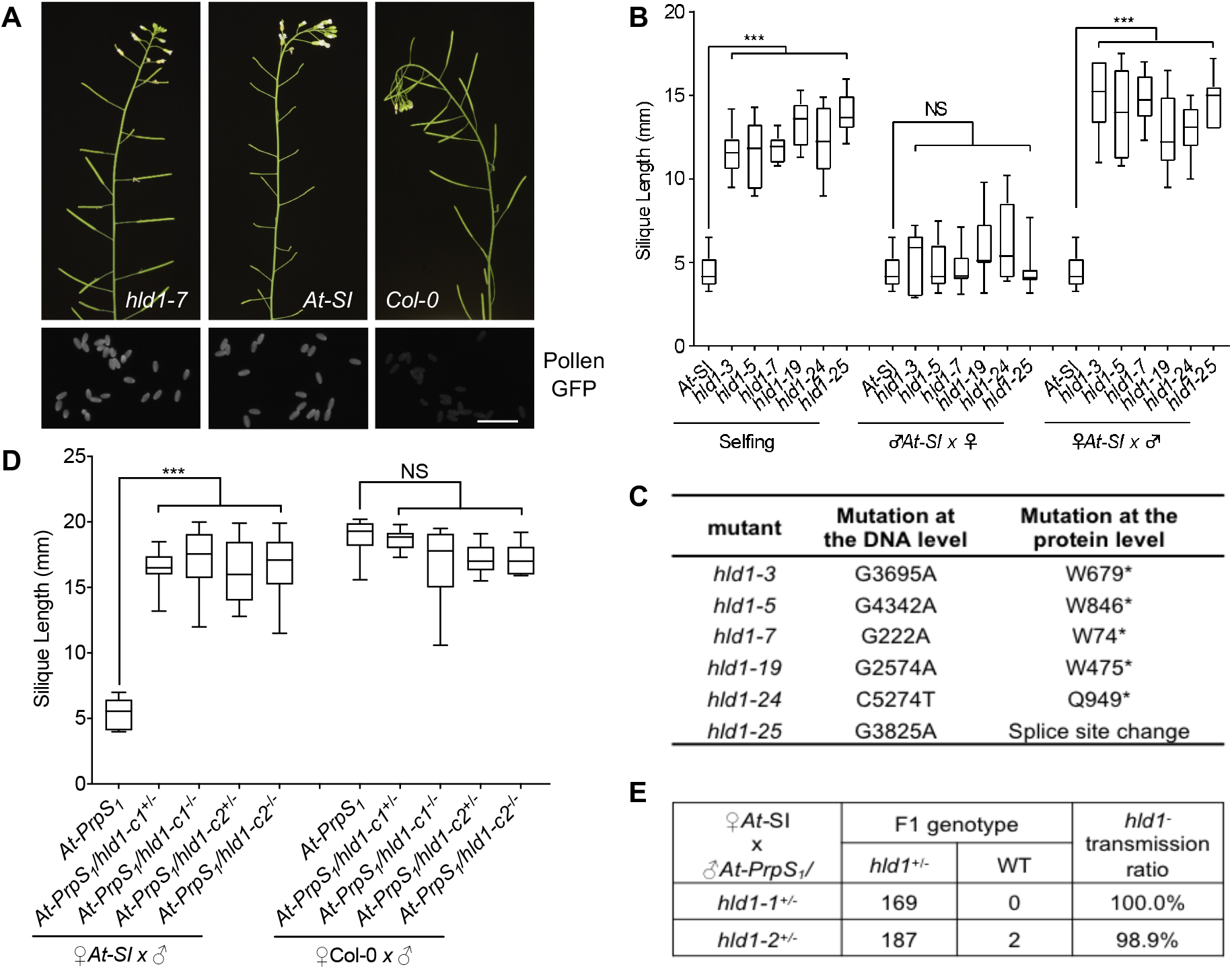
Identification of *hld1* as a male gametophytic mutant that overcomes SI. (A) ***hld1* was identified from an EMS mutagenesis screen**. Mutants were screened for a defective SI phenotype, with normal seed set. Upper panel: Inflorescences from *hld1-7* mutants had normal seed set like WT *Col-0* plants, in contrast to *At-*SI. Lower panel: PrpS_1_-GFP signals in *hld1* mutant pollen are similar to those in *At-*SI pollen. Bar = 100 μm. (B) ***hld1* mutants have pollen defects**. *hld1* stigmas were pollinated with self-pollen (selfing) or *At-*SI pollen (♂At-*SI* x ♀) and *At-*SI stigmas pollinated with *hld1* pollen (♀At-*SI* x ♂). *At-* SI stigmas pollinated with *At-*SI pollen acted as the SI controls for these pollinations. Silique length measurements (B) and number of seeds per silique (Fig S2C) showed that the *hld1* mutants used as the male parent had significantly longer silique lengths and higher seed-set than the *At-*SI parent line. *hld1* mutants used as the female parent pollinated with *At*-SI pollen displayed a normal SI phenotype of short silique lengths and low seed set. N=9-11. One-way ANOVA. ***: p<0.001. NS: not-significant, p>0.05. (C) **Six different *At3g27325* mutant alleles were identified for the *hld1* mutants through WGS**. * indicates stop codon. (D) **Mutation of *At3g27325* phenocopies *hld1***. *At-*SI or Col-0 stigmas were pollinated with *At-PrpS*_*1*_ pollen containing the CRISPR/Cas9-derived *At3g27325* mutant allele (*hld1-c1* or *hld1-c2*). Significant increases in both the silique lengths (C) and seed set (Fig S5A) were observed when *At3g27325* was mutated, in both heterozygous and homozygous mutants. N=12-16. One-way ANOVA. ***: p<0.001. NS: not-significant, p>0.05. (E) ***hld1* mutants are male gametophytic mutants**. Segregation analysis of the F1 population of ♀*At*-SIx♂*At-PrpS*_*1*_*/hld1*^*+/-*^ showed that ∼100% of the progenies are *hld1* heterozygous mutants, demonstrating that when pollinated on *At*-SI stigmas, only *hld*^*-*^ mutant pollen can bypass the SI response, resulting in successful fertilization and seed set.

### WGS identified *At3g27325* as the causal gene for all the *hld* mutants

To reveal the molecular identity of the *hld* mutants, we used whole genome sequencing (WGS) followed by SNP analysis to map the causal gene of these *hld* mutants. We sequenced the genome of a self-compatible population of six independent mutants after two backcrosses. SHOREmap analysis^21^ revealed six different nonsynonymous mutations in a single gene, *At3g27325* (**Fig 1C, Fig S3**). Analysis of the segregating populations revealed 100% linkage between the *At3g27325* mutation and suppression of SI (**Table S2**, n=298). Targeted genotyping of the remaining *hld* mutants identified six additional independent mutant alleles of the same gene (**Table S3**). These results indicate that our screen reached a high level of saturation and suggest that *At3g27325* (designated as *HLD1* hereafter), is the single causal locus for SI suppression in all the mutants identified here.

### CRISPR/Cas9 confirms that *At3g27325* is the causal gene of *hld1* mutant phenotype

To confirm *At3g27325* as the causal gene for SI suppression, we generated two independent CRISPR knockout mutants (*hld1-c1* and *hld1-c2*) with Arabidopsis transgenic line *pNTP303::PrpS*_*1*_*-GFP* (*At-PrpS*_*1*_) as the background (**Fig S4**), as the non-mutagenized parent line *At*-SI is self-incompatible and unfeasible to use. Pollinating *At-*SI stigmas expressing *PrsS*_*1*_ with *At-PrpS*_*1*_ pollen resulted in a SI phenotype with short siliques and no seed set. However, *At-*SI stigmas pollinated with *At-PrpS*_*1*_/*hld1-c1* or *At-PrpS*_*1*_/*hld1-c2* pollen displayed normal siliques and seed set (**Fig 1D, Fig S5A**). This demonstrates that targeted knockout mutation of *At3g27325* is sufficient to prevent SI. Consistent with this, the F1 population of an *At-*SI x *At-PrpS*_*1*_*/hld1-c1/2*^*+/-*^ cross consisted of 99.5% *hld1*^*+/-*^ heterozygous plants (n=358; **Fig 1E**), showing that it is almost exclusively *hld1* mutant pollen that transmits its genome to the next generation. When introgressed into the *At*-SI background line, *hld1-c1* and *hld1-c2* were both sufficient to cause a breakdown of SI upon self-pollination, comparable to those EMS mutagenesis-derived *hld1* alleles (**Figure S5B, C**). Our finding that CRISPR-generated mutations of *At3g27325* phenocopy the EMS-generated *hld1* mutant phenotype confirms that *At3g27325* is the causal gene for *hld1*-mediated SI suppression.

### SI-suppression by *hld1* does not depend on specific *S*-alleles

Successful prevention of self-pollination by SI relies on a polymorphic *S-*locus involving *S*-specific self-recognition. Although the data presented so far show that *HLD1* regulates the *PrpS*_*1*_*-PrsS*_*1*_-based SI response, it was not clear if *HLD1* only regulates *PrpS*_*1*_*-PrsS*_*1*_-based SI, or if it acts as a genuine *S*-specific *Papaver* SI regulator. To examine if the *hld1* mutants regulate SI independent of specific *S*-alleles, we examined if the *hld1* mutation disrupted SI in plants expressing an alternative pair of cognate *S*-alleles, *PrpS*_*3*_ and *PrsS*_*3*_. We introduced three *hld1* mutant alleles, *hld1-c1, hld1-c2*, and *hld1-7*, into a *pNTP303::PrpS*_*3*_-*GFP* line (*At-PrpS*_*3*_). Consistent with previous observations, when *Col-0* stigmas were pollinated with *Col-0*, or *At-PrpS*_*3*_, or *At-PrpS*_*3*_*/hld1* pollen, there was no significant difference observed in the silique length and number of seeds per silique^19^. Pollinating plants expressing the *PrsS*_*3*_ in the stigma (*At-PrsS*_*3*_) with *At-PrpS*_*3*_ pollen resulted in SI, with a significant reduction in silique length and almost no seed, compared with the control pollination ♀*At-PrsS*_*3*_ x ♂*Col-0* (**Fig 2A, Fig S6**). In contrast, pollinating *At-PrsS*_*3*_ with *At-PrpS*_*3*_*/hld1* pollen resulted in normal silique lengths and seed set (**Fig 2A, Fig S6**), demonstrating that mutation of the *HLD1* gene abolishes the *PrpS*_*3*_*-PrsS*_*3*_-based SI. These results demonstrate that *HLD1* regulates SI irrespective of the *S*-alleles involved.

**Fig. 2.**
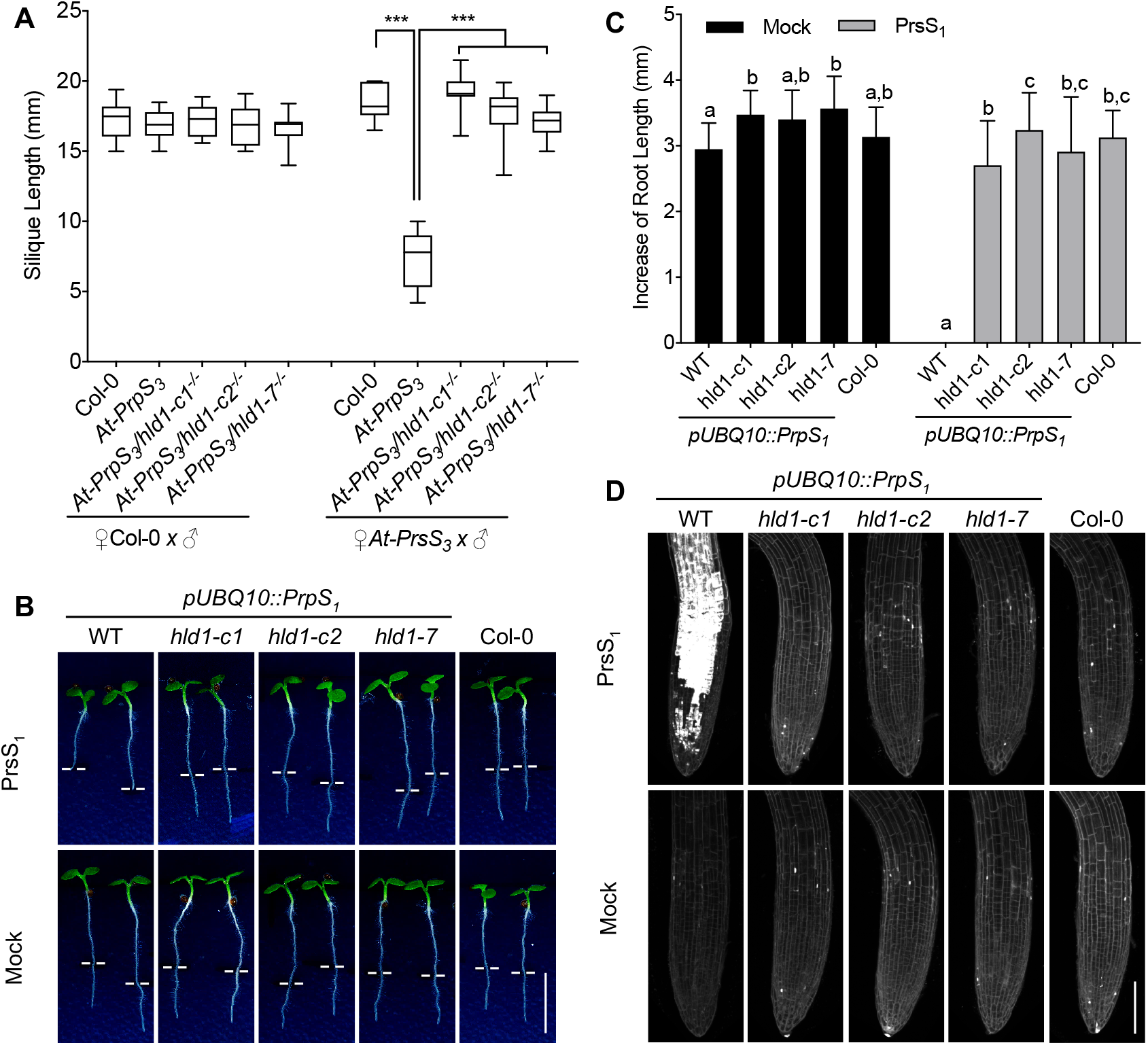
*HLD1* regulates SI in an *S*-specific manner and regulates ectopic “SI” in roots. (A) ***HLD1* regulates SI in an *S*-specific manner**. *At-PrpS*_*3*_ pollen containing *hld1-c1, hld1-c2, or hld1-7* mutant allele were pollinated onto *At-PrsS*_*3*_ or Col-0 stigmas. Silique lengths (A) and seed set (Fig S6) were measured. N=10-20. One-way ANOVA. ***: p<0.001. (B-D) ***HLD1* regulates ectopic “SI” in Arabidopsis roots**. (B) Treatment of *pUBQ10::PrpS*_*1*_ seedlings with PrsS_1_ resulted in root growth inhibition; in contrast, when *pUBQ10::PrpS*_*1*_*/hld1* seedlings were given the same treatment, no inhibition was observed (upper panel). Mock treatment did not affect any of the seedling roots (lower panel). White lines indicate the position of root tips when treated. Bar = 0.5 cm. The increases of root length 24h after treatment were quantified in (C). 12-20 seedlings from three independent experiments were documented for each treatment. Results = mean ± SD. One-way ANOVA with multiple comparison test. Different letters indicate p<0.05. (D) Treatment of *pUBQ10::PrpS*_*1*_ seedlings with PrsS_1_ resulted in root cell death indicated by PI signals (white). However, when *pUBQ10::PrpS*_*1*_*/hld1* seedlings were treated with PrsS_1_ proteins, no cell death was observed, comparable with what was observed for *Col-0* seedlings. Bar = 100 μm.

### HLD1 regulates ectopic “SI” in Arabidopsis roots

We recently established that expression of *PrpS*_*1*_*-PrsS*_*1*_ in Arabidopsis roots triggers an ectopic “SI-like PCD” response in vegetative tissues, resulting in root growth inhibition and death of root cells^22^. To test if *HLD1* is necessary for this ectopic “SI-like PCD” response outside the reproductive context, we introduced *hld1-c1, hld1-c2* and *hld1-7* into the *pUBQ10::PrpS*_*1*_ transgenic line. While addition of recombinant PrsS_1_ protein to plants carrying *pUBQ10::PrpS*_*1*_ led to root growth arrest and cell death^22^, treatment of *pUBQ10::PrpS*_*1*_*/hld1* seedling roots with PrsS_1_ showed no such effect (**Fig 2B-D**). This suppression of root growth inhibition and root cell death reveals that *HLD1* is not only required for SI, but also for the ectopic “SI-like PCD” response independent of the reproductive context.

### *HLD1* is a *HsPGAP1* ortholog

*HLD1* contains a coding region of 3255 bp encoding a predicted protein of 1085 amino acids. Protein sequence analysis revealed that HLD1 contains a PGAP1-like domain (PFAM domain ID: PF07819), and is a putative PGAP1 homologue^6,14^. *PGAP1* has been identified in mammals and its orthologue, *<u>B</u>ypass of <u>S</u>ec <u>T</u>hirteen 1* (*Bst1*) in yeast; it encodes a GPI inositol-deacylase. However, although the databases indicate the presence of *PGAP1*-like gene in plants, no analysis of *PGAP1* or the function of the protein it encodes has been made in plants to date.

To understand the evolution of the PGAP1 proteins, we constructed a phylogenetic tree of 631 predicted PGAP1 protein homologues from > 300 eukaryotic species (**Fig S7A, B**). Two major phylogenetic clades were identified for plants; in Arabidopsis, HLD1 and another PGAP1-like domain containing protein At5g17670 were classified into two different clades (**Fig S7A**). The divergence of these two clades can be traced back to an ancient whole genome duplication event occurring before the radiation of extant Viridiplantae, including green algae and the land plants (**Fig S7A, B**)^23^. Comparison of the two homologous, HLD1 and At5g17670 from *A. thaliana* with *Homo sapiens* PGAP1, revealed that HLD1 has a slightly higher amino acid sequence similarity to *Hs*PGAP1 (23.3%) than At5g17670 (20.1%). Importantly, HLD1 shares a similar secondary protein structure to *Hs*PGAP1 while At5g17670 does not (**Fig S7C**). This suggests that HLD1 is likely to be the *Hs*PGAP1 ortholog in Arabidopsis^6,14^.

### HLD1 functions as a GPI-inositol deacylase

Human PGAP1 is a GPI-inositol deacylase that removes the inositol acyl chain after the attachment of the GPI anchor to its target protein^12^. *Hs*PGAP1 is ubiquitously expressed in humans^24^, and likewise we found HLD1 expression in all the Arabidopsis tissues examined (**Fig S8, S9**)^25^. A key feature of PGAP1 is a catalytic serine-containing motif, V***GHSMGG, in the PGAP1-like domain that is highly conserved across the eukaryotic kingdoms. The serine 218, at the active site is also present in *HLD1* (**Figure 3A**). Mutation of *PGAP1* in mammals results in a “three-footed” structure of GPI-APs that retain an extra acyl chain in addition to the two membrane-anchored non-polar fatty acid tails^12^. This particular conformation makes GPI-APs in the plasma membrane insensitive to cleavage-mediated release from the plasma membrane by phosphoinositide-specific phospholipase C (PI-PLC)^12^. We therefore attempted to establish the biochemical function of the Arabidopsis HLD1 by examining if a GPI-AP in the *hld1* mutant background contained the third inositol acyl chain and was resistant to PI-PLC treatment. To verify that HLD1 functions as a GPI-inositol deacylase, we introduced the *hld1* mutant into a line expressing GFP-tagged SKU5, which was adopted as a representative Arabidopsis GPI-AP localizing to the plasma membrane and cell wall^26^. A PI-PLC assay revealed that GFP-SKU5 was highly resistant to PI-PLC-mediated GPI cleavage and was not released from plasma membrane in the *hld1* mutant (**Fig 3B**). This result recapitulates analogous results obtained in mammalian *pgap1* knockouts^9,12,27^ and supports the idea that *GFP-SKU5/hld1* has three tails in the GPI anchor and is therefore insensitive to PI-PLC treatment^12^, providing good evidence that HDL1 is an inositol deacylase and a functional ortholog of PGAP1.

**Fig. 3.**
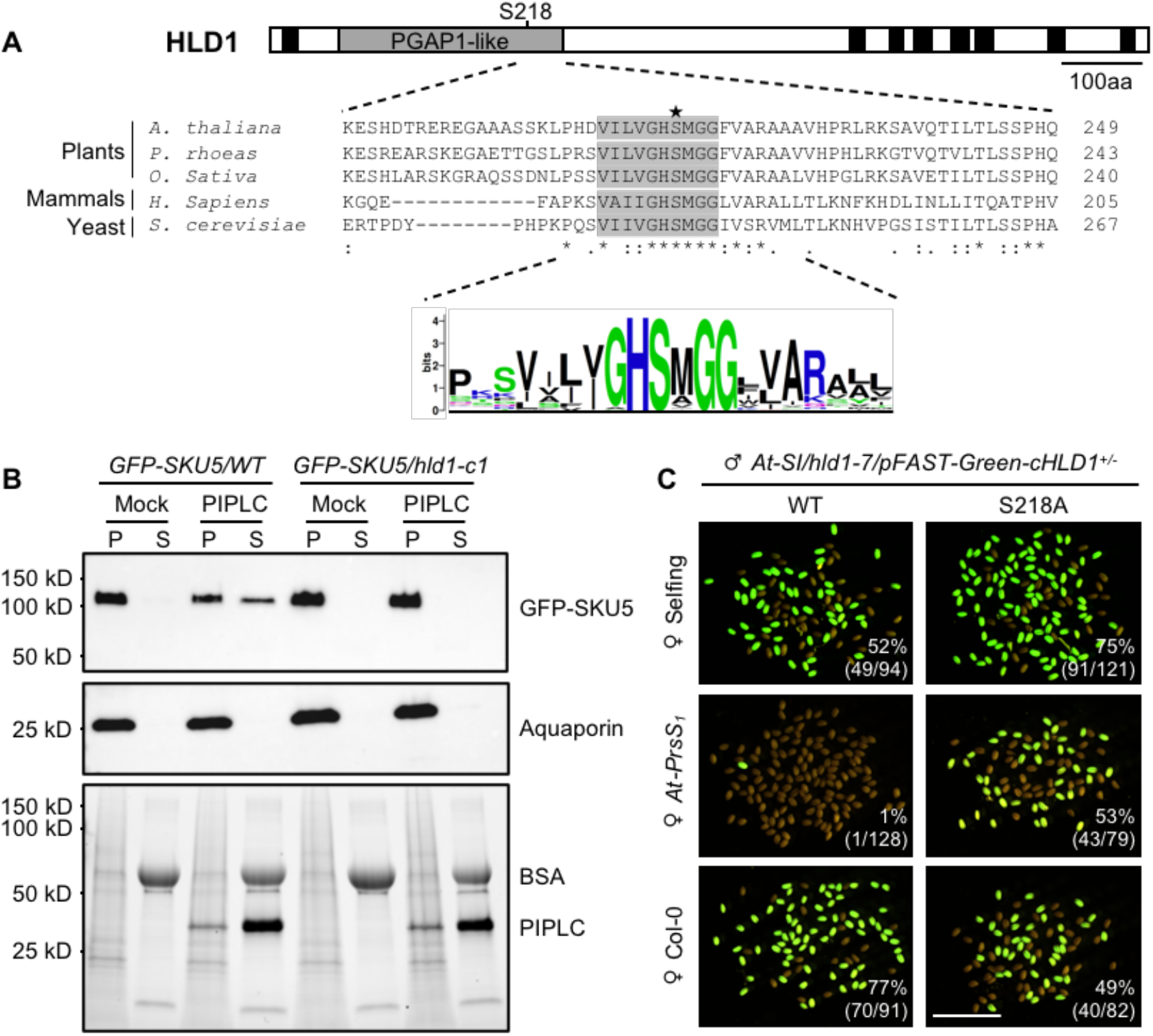
HLD1 is a GPI inositol deacylase and its lipase activity is required for the SI response. (A) **The lipase motif in the PGAP1-like domain is highly conserved across the eukaryotic kingdoms**. Upper panel: cartoon of HLD1 protein secondary structure. The PGAP1-like domain is indicated by the grey box. Black boxes indicate transmembrane domains. Middle panel: amino acid sequence alignment around the conserved lipase motif in the PGAP1-like domain in several higher plants, human and yeast. The grey box indicates the lipase motif with the catalytic site Serine 218 indicated (star). Bottom panel: amino acid motif logo shows that the lipase motif of the PGAP1-like domain is conserved across eukaryotic kingdoms. (B) **HLD1 functions as a GPI inositol deacylase**. GFP-SKU5 was enriched in the pellet (P) fractions in both wildtype (WT) and *hld1* mutants in the mock (buffer control) treatment. PI-PLC treatment resulted in the GFP-SKU5 being found in the soluble (S) fraction in WT samples, showing cleavage by PI-PLC, while in the *hld1* mutants, GFP-SKU5 remained in the pellet, demonstrating that in the *hld1* mutant this GPI-AP was resistant to PI-PLC treatment. This supports the idea that in the *hld1* mutant a persistent inositol-linked acyl chain makes them insensitive to cleavage by PI-PLC. (C) **HLD1 lipase activity is required for the SI response**. *Hld1* mutants were transformed with *pHLD1::mCherry-cHLD1* or *pHLD1::mCherry-cHLD1(S218A)* cloned in the pFAST-Green plasmid vector backbone. Representative images of seeds resulting from pollinations by pollen from T1 heterozygous plants onto self-stigmas (upper panel), *At-PrsS*_*1*_ stigmas (middle panel) or *Col-0* stigmas (lower panel). GFP signals report the pollen transmission in the seeds in the F1 generation. Numbers indicate the ratio of GFP-positive seeds (number of GFP seeds/total seeds). Bar = 5 mm.

### *HLD1* GPI-inositol deacylase function is required for SI

To examine if HLD1 inositol deacylase activity is required for the SI response, we mutated the conserved serine 218 in the catalytic core of the PGAP1-like domain^12^ (**Fig 3A**) to alanine (S218A), and cloned it into the pFAST-Green plasmid vector backbone^28^. The use of pFAST-Green facilitates the examination of transmission of gametes by simply checking the green fluorescence of the seeds. In segregating T2 seeds from heterozygous *At-*SI*/hld1/cHLD1* plants, only around 50% of the seeds were GFP positive. In contrast, heterozygous *At-* SI*/hld1/cHLD1(S218A)* plants displayed normal Mendelian segregation, with 75% GFP-positive T2 seeds, as expected for a full transmission via both germlines (**Fig 3C; Table S4**). This suggests that pollen containing the *cHLD1* transgene did not achieve fertilization and provides evidence for the failure of the S218A mutant allele to restore the SI response. This hypothesis was verified by the observation that almost no GFP positive seeds were found when *At-SI/hld1/cHLD1* pollen was pollinated onto *At-PrsS*_*1*_ stigmas, which was not observed for *At-SI/hld1/cHLD1(S218A)* pollen. Thus, unlike the wild-type *HLD1* that rescued the *hld1* phenotype and restored SI, the inositol deacylase-defective S218A *HLD1* allele did not (**Fig 3C; Table S4**). This demonstrates that the HLD1 inositol deacylase activity is essential for SI, implicating a requirement for correctly remodeled GPI-APs in this process.

### GPI-APs at the membrane is required for SI

To test this hypothesis, we next examined if the absence of GPI-APs at the plasma membrane had a similar effect on SI as *hld1. SETH1* and *SETH2* are orthologs of *PIG-C* and *PIG-A* genes involved in early GPI biosynthesis, respectively; knockouts lack expression of GPI-APs at the plasma membrane^5,27^. We introduced *seth1-2* and *seth2* mutant alleles^10^ into the *At-PrpS*_*1*_ background. As loss of *SETH1* or *SETH2* function results in homozygote lethality^10^, we generated heterozygous *At-PrpS*_*1*_*/seth1-2*^*+/-*^ and *At-PrpS*_*1*_*/seth2*^*+/-*^ lines. Pollinating *At-PrsS*_*1*_ stigmas with pollen from *At-PrpS*_*1*_*/seth1-2*^*+/-*^ or *At-PrpS*_*1*_*/seth2*^*+/-*^ plants produced short siliques without seeds, suggesting initially that mutations in these genes did not prevent the SI phenotype (**Fig 4A, Fig S10A**). However, consistent with previous report, *seth1-2* and *seth2* showed almost completely abolish pollen germination and tube growth (**Fig S10B**)^10^, thus potentially masking the role of these genes in SI. We therefore used the *in-vitro* SI bioassay^29^ with ungerminated pollen to examine the effect of these mutants on pollen viability and cell death. Addition of recombinant PrsS_1_ to *At-PrpS*_*1*_*/seth1-2*^*+/-*^ or *At-PrpS*_*1*_*/seth2*^*+/-*^ pollen, resulted in close to 50% pollen grain death in contrast to > 98% death in *At-PrpS*_*1*_ pollen (**Fig 4B, C, Fig S10C**). This demonstrates that *SETH1* or *SETH2* knockouts prevent SI-induced cell death and implicates the presence of GPI-APs at the plasma membrane being indispensable for SI-PCD.

**Fig. 4.**
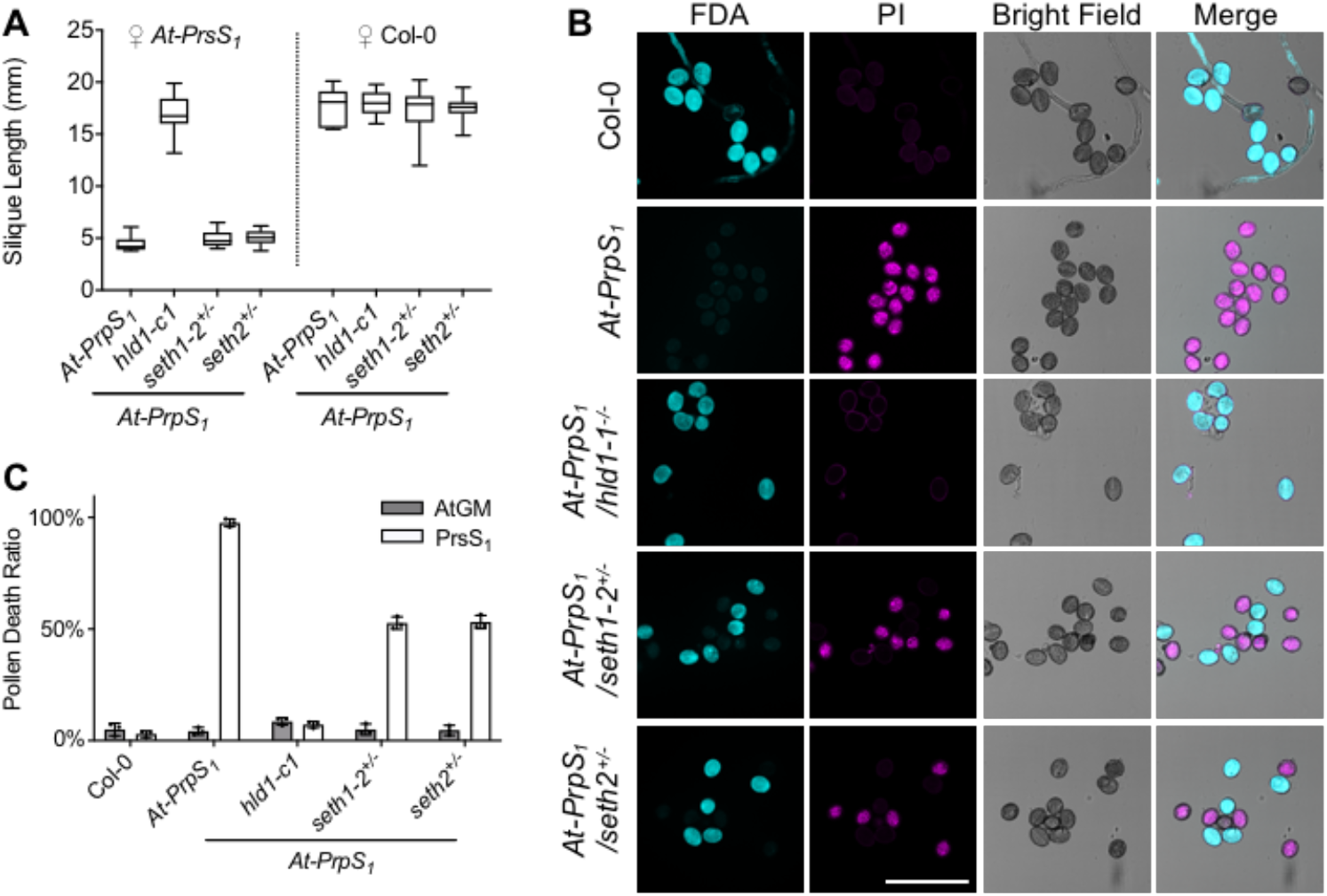
The GPI-anchoring pathway is involved in the regulation of SI-induced death of pollen. (A) **Mutation of GPI-anchoring pathway genes does not phenocopy the hld1 mutant phenotype**. GPI-anchoring pathway mutants, *seth1-2* and *seth2*, were introduced into *At-PrpS*_*1*_ background, and the pollen were pollinated onto *At-PrsS*_*1*_ or Col-0 stigmas. Pollinating *At-PrsS*_*1*_ stigmas with *At-PrpS*_*1*_*/seth1*^*+/-*^ or *At-PrpS*_*1*_*/seth2*^*+/-*^ pollen resulted in short siliques (A) and no seedset (Fig S10A). This is in consistent with the observation that *seth1-2* and *seth2* abolished the pollen fertilization ability, showing almost no transmission (Fig S10B). N=13-17. (B-C) **Mutation of GPI-anchoring pathway genes resulted in the alleviation of SI-induced PCD**. *At-PrpS*_*1*_*/seth1*^*+/-*^ and *At-PrpS*_*1*_*/seth2*^*+/-*^ pollen were treated with PrsS_1_ proteins (B) or Mock buffer control (Fig S10C). The samples were co-stained with FDA and PI 6h after treatment. Bar = 100 μm. The PI positive ratios were quantified in (C). Three independent experiments were carried out. In each experiment, 100-200 pollen grains were counted for each of the sample.

### *HLD1* mutation does not affect the expression levels of GPI-APs

In mammals and yeast, GPI-inositol deacylation by PGAP1 homologs^12^ is important for efficient sorting of GPI-APs to exit the ER^5,30^. However, knockout of *PGAP1* does not prevent transport of GPI-APs to the plasma membrane, so the precise functional significance of GPI deacylation remains unclear^31^. To examine if prevention of GPI inositol deacylation in the *hld1* mutant also fails to affect the expression of GPI-APs at the plasma membrane in plants, we examined GFP-SKU5 levels (**Fig S11A**). Imaging of GFP-SKU5 confirmed that the *HLD1* mutation had no obvious effect on GFP-SKU5 localization at the plasma membrane (**Fig 5A; Fig S11B**). Quantitation of the GFP-SKU5 signals revealed that the GFP-SKU5 protein levels in *hld1* mutant seedlings were not significantly different from the wild type (p=0.7995, n= 8; **Fig 5B**). This is consistent with the results of *PGAP1* knockout in animals^12,27^ and is in line with our observation that the *hld1* mutants had no obvious effect on overall Arabidopsis development, except for slightly delayed onset of flowering (**Fig S12**). This contrasts with the homozygous *seth1/2* mutants, which are lethal, and *gpi8-1* mutants, which show reduced GPI-APs expression and disrupted plant growth (**Fig S12**)^32^. These results, demonstrating that *hld1* does not affect GPI-AP abundance at the plasma membrane, show that *hld1* is not likely to affect SI due to failure of GPI-AP targeting to the plasma membrane, implying a role of GPI-AP remodeling in SI.

**Fig. 5.**
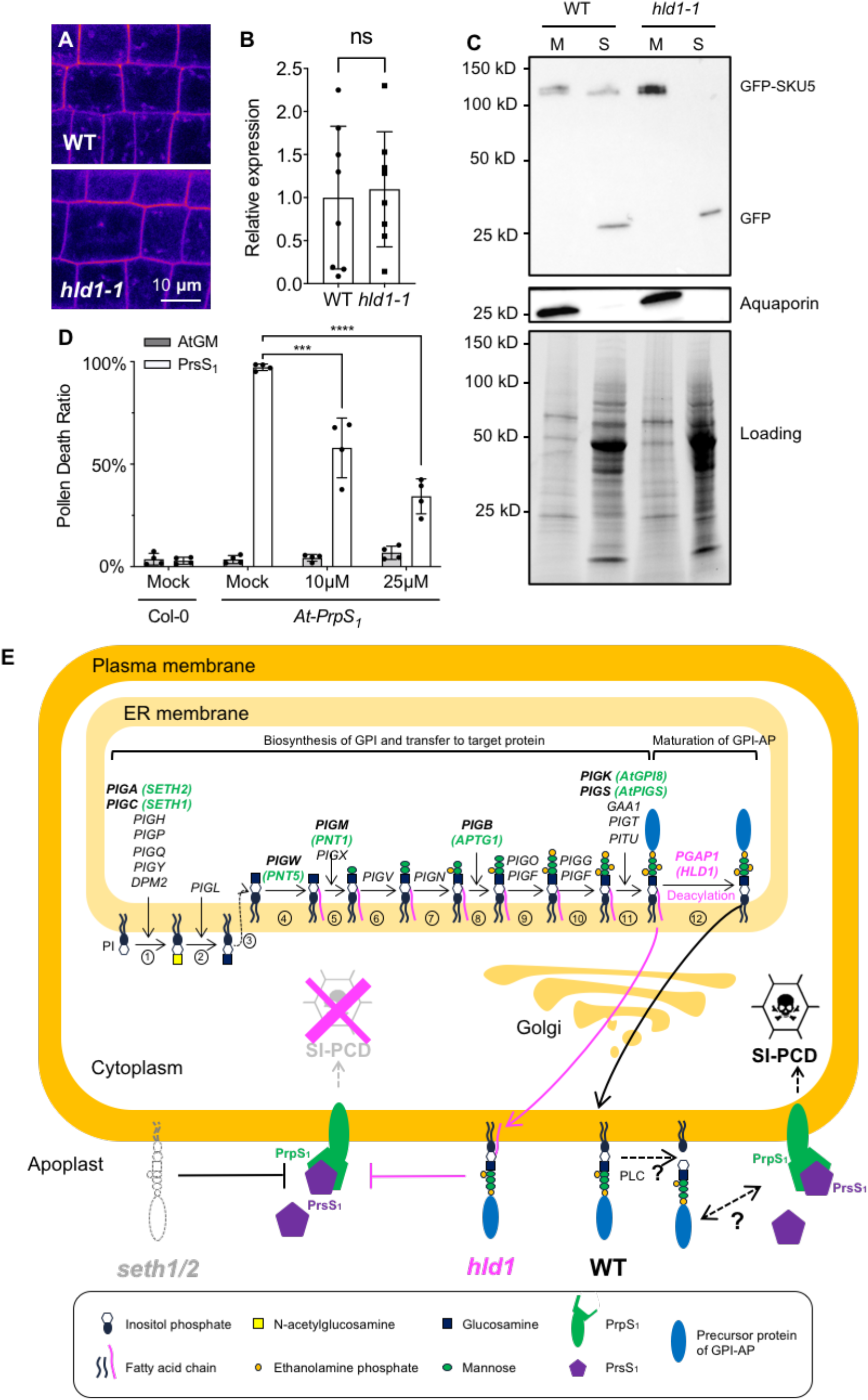
HLD1 regulates SI by affecting PLC-mediated release of GPI-APs. (A-B) **Mutation of HLD1 does not affect GFP-SKU5 protein expression level and localization**. (A) Representative confocal images (single Z slice) of GFP-SKU5 signals of 4-day-old *GFP-SKU5/WT* or *GFP-SKU5/hld1-1* seedlings. (B) Quantitation of the protein expression level of GFP-SKU5 using western blot analysis of F3 seedlings derived from 16 F2 double homozygous siblings of the cross ♀*hld1-1* x ♂*GFP-SKU5*. Tubulin was used as an internal control. Quantification revealed that GFP-SKU5 protein expression level was not altered by the HLD1 mutation (p=0.7995, Students t test). These data show that this GPI-AP was still targeted to the plasma membrane in the *hld1-1* mutant plants that lack inositol deacylase activity. (C) **The HLD1 mutation inhibits the cleavage and release of GFP-SKU5 from the plasma membrane**. GFP-SKU5/WT and GFP-SKU5/hld1 proteins were extracted, separated into membrane (M) and soluble, supernatant (S) fractions, and analyzed using western blots. In the WT samples, GFP-SKU5 was detected in both membrane and soluble fractions; in contrast, the *hld1* mutant, GFP-SKU5 was only observed in the membrane fraction. This provides evidence that cleavage and release of GPI-APs from the plasma membrane is prevented in the *hld1-1* mutant. (D) **The PLC inhibitor U73122 treatment inhibits SI-induced pollen death**. *Col-0* and *At-PrpS*_*1*_ pollen that were “mock treated” with Arabidopsis pollen germination medium (AtGM) exhibited low levels of death; *At-PrpS*_*1*_ pollen treated with recombinant PrsS_1_ had high levels of death, and U73122 alleviated death in a dose-dependent manner. This suggests that cleavage of GPI-APs by PLC is an important requirement for mediating SI-induced death of pollen. (E) **Cartoon of key events in GPI-AP biosynthesis/maturation and how *pgap1*/*hld1* affects self-incompatibility**. GPI-anchoring is a post-translational modification involving several phases which have been established in mammals and yeast. First, biosynthesis of the GPI involves a series of events [steps 1-10]. Once the core GPI is assembled [step 10], the target precursor protein (blue sphere) is transferred to the GPI by a GPI transamidase complex [step 11]. The nascent immature GPI-AP then undergoes remodeling; the first of these is deacylation [step 12], involving PGAP1, which we identified in this study as HLD1 (indicated in pink). After this several other PGAP genes (not shown) mediate further modification during the transport from ER to the Golgi and then to the plasma membrane. The genes involved identified in humans are indicated in black; genes identified in plants are indicated in green. Knockouts of genes early in the biosynthetic pathway in mammals result in lack of expression of GPI-APs at the plasma membrane^5,27^; in plants homozygous knockouts of the orthologs of *PIG-C* and *PIG-A, SETH1* and *SETH2*, are lethal. Our data show that correctly modified GPI-APs at the plasma membrane are crucial for the *Papaver* SI response. This SI response is controlled by interaction of membrane-bound PrpS and secreted PrsS. Interaction of cognate PrpS-PrsS triggers the SI-response leads to growth arrest and programmed cell death (PCD) specifically of “self”-pollen. This SI-PCD response is prevented in GPI biosynthesis and GPI-remodeling mutants. When interfering with GPI biosynthesis by SETH1 or SETH2 mutation no SI-PCD occurs, suggesting that presence of GPI-APs at the plasma membrane are required for SI. The GPI-remodeler HLD1 removes the acyl chain linked to the GPI in nascent GPI-APs [step 12]. In the *hld1* mutant, this step does not occur. As a consequence, the GPI-APs retain their inositol-linked acyl chain (indicated in pink), resulting in a “3-footed” configuration that retains the GPI-AP in the plasma membrane. We show that knockout of HLD1 completely prevents SI. This demonstrates that GPI-AP remodeling by inositol deacylation is critical for the SI response. As release of GPI-APs from the plasma membrane by phospholipase C activity (PLC) is implicated, it is tempting to speculate that release of GPI-APs into the apoplast is critical for SI. Based on these observations, we propose that SI depends on direct or indirect interaction of released apoplastic GPI-APs. *Cartoon adapted from*^*9*^.

### Inositol deacylation by HLD1 is required for release of GPI-APs

Besides their localization in the outer leaflet of the plasma membrane, many GPI-APs in mammals can be cleaved and released into the extracellular space; this can be mediated by phospholipase C (PLC) and is important for many cellular processes, including adhesion, proliferation, survival and oncogenesis^9,13^. In wild-type Arabidopsis, we detected GFP-SKU5 in both the membrane and the soluble fraction (**Fig 5C**); this is consistent with a report that SKU5 can be cleaved and released into the apoplast^26^. In contrast, in the *hld1* mutant, GFP-SKU5 was exclusively detected in the membrane fraction (**Fig 5C**). Moreover, using a Triton X-114 assay, GFP-SKU5 was detected primarily in the aqueous phase in wild-type, whereas the majority of GFP-SKU5 was present in the detergent phase in the *hld1* mutant (**Fig S13**). These results demonstrate that in absence of HLD1, GFP-SKU5 is more hydrophobic and is retained at the plasma membrane. This provides evidence that release of GFP-SKU5 (and probably other GPI-APs) from the plasma membrane requires GPI inositol deacylation by HLD1.

### HLD1 regulates SI by affecting PLC-mediated release of GPI-APs

As release of the GPI-AP SKU5 from the plasma membrane by GPI cleavage is prevented by the “three-footed” configuration in *PGAP1* mutants^9,12,27^, it is conceivable that the prevention of SI by the *HLD1* mutation is caused by a failure to release GPI-APs from the plasma membrane into the apoplast. Studies on PGAP1 using animal cell lines lacking a GPI inositol deacylase showed that GPI-APs were resistant to PLC-mediated release, showing that this release can be mediated by PLC^12^. We therefore investigated this possibility further by testing whether inhibition of PLC activity might prevent SI in *At-*SI pollen. We applied three different phospholipase C inhibitors (U73122, ET-18-OCH3, and C48/80)^33,34^ to the pollen *in vitro* SI-bioassay. As pollen tube growth requires PLC activity^35,36^, we could not test the effect of these inhibitors to see if they prevent SI-induced pollen tube growth inhibition, so we examined if the PLC inhibitors prevented the SI-induced death of pollen. *At-PrpS*_*1*_ pollen pretreated with the PLC inhibitor U73122 significantly alleviated SI-induced cell death in a dose-dependent manner, with death reduced by ∼40% at 10 μM (p<0.001), and by ∼63% at 25 μM U73122 (**Fig 5D**, p<0.001). Pretreatment with the other two PLC inhibitors, ET-18-OCH3, and C48/80, showed similar alleviation of SI-induced pollen death in a dose-dependent manner (**Fig S14**).

This reduction in death rates by PLC inhibitor treatment demonstrates that PLC activity is important for SI-induced pollen death, suggesting an important role of PLC-mediated release of GPI-APs from the plasma membrane in the SI response. As SI is prevented by the “three-footed” GPI-AP configuration resulting from the *HLD1* mutation, our evidence is not only consistent with the idea that GPI-APs play a key role in SI, but that specifically the prevention of their release from the plasma membrane is responsible for the breakdown of SI in the *hld1* mutant plants.

## Discussion

We identified *HLD1*, a previously uncharacterized plant *PGAP1* ortholog that is essential for the *Papaver* SI-response. We provide evidence that GPI anchoring, and specifically GPI deacylation of GPI-APs, is essential for the SI response. Although a putative *PGAP1* ortholog has been identified in the Arabidopsis genome databases^6,14^, our study provides the first functional analysis of a GPI-remodeling protein in plants. In animals, rather little is known about the physiological functions of PGAP1 apart from its biochemical role in deacylation in the GPI-anchoring pathway^12^ and pathological consequences of PGAP1 mutations^27,31,37,38^. *PGAP1* knock-out mice display serious developmental defects^31^ and humans with a null mutation in *PGAP1* suffer from intellectual disabilities and encephalopathy^27,31,37,38^. This shows that retention of an extra acyl chain in GPI anchors causes severe developmental defects in mammals. In contrast, in plants lack of GPI deacylation does not produce grave developmental defects, but specifically interferes with the SI process in *At*-SI lines. As it has been proposed that the lack of GPI deacylation disturbs GPI anchor-mediated signal transduction^31^, prevention of deacylation in the *hld1* mutants may affect cell-cell signaling involved in SI. Intriguingly, our data suggest that the cleavage and membrane release of GPI-APs may play an important functional role in this context. As release of GPI-APs from the plasma membrane by GPI cleavage is prevented by the “three-footed” configuration in *PGAP1* mutants^9,12,27^, and as we showed that PLC inhibitors alleviate SI-induced death, our data are consistent with the idea that prevention of SI by the *HLD1* mutation is caused by a failure to release GPI-APs from the plasma membrane into the apoplast. This suggests an important role for the release of GPI-APs from the plasma membrane in this SI response.

Our identification of *HLD1* implicates an involvement of GPI-APs in the SI response. As *seth* mutants have the same detrimental effect on SI as *hld1* mutants, the presence of GPI-APs at the plasma membrane appears to be crucial (**Fig 5E**); While PrpS itself is clearly not a GPI-AP, so cannot be a substrate of HLD1^39^, PrpS functions might be regulated by GPI-APs (**Fig 5E**). In mammalian cells, GPI-APs are involved in key events such as embryogenesis, development, and fertilization^8^. Over 250 GPI-APs have been predicted in Arabidopsis^4,40^; though despite recent advances in our knowledge of GPI-APs in plants, information on their functional roles remains limited. There is currently intense interest in plant GPI-APs, as they play a key role in cell-cell signalling, through association with partner receptor-like kinases^41-43^. It appears that broadly expressed GPI-APs, rather than those specifically involved in reproduction, are involved in SI, as the ectopic “SI-like PCD” response in Arabidopsis roots also requires *HLD1*. Our study opens up new avenues for research into the involvement and regulation of GPI anchoring in SI, investigation of the GPI-anchoring pathway and the identity and roles of GPI-APs in plant signalling processes in general.

## Acknowledgements

We thank members of the PCD laboratory for critical comments on the manuscript, and Ms. Freya De Winter for technical assistance. This research was financially supported by the European Research Council (ERC) StG PROCELLDEATH 639234 and CoG EXECUT.ER 864952 (MKN), Biotechnology, Biological Sciences Research Council (BBSRC) grant BB/P005489/1 (VEF-T, MB), Fonds Wetenschappelijk Onderzoek (FWO) grant G011215N (MT), FWO grant 12I7417N (ZL), Chinese Scholarship Council (CSC) grant 201806760049 (FX).

## Author Contributions

ZL, MKN, VEF-T, and MB designed the study. ZL, FX, and MT performed the research and analyzed data. ZT contributed to the PGAP1 phylogeny analysis. FC and LS contributed to the analysis of WGS data. ZL, VEF-T, MKN and MB wrote the manuscript with input from all the other authors.

## Data and materials availability

All data generated or analyzed during this study are included in this published article (and its supplementary information files). All the materials used in this study will be freely available to the scientific community upon request under MTA.

## Accession numbers

Sequence data from this article can be found in the GenBank/EMBL data libraries under accession numbers: *AtGPI8(At1g08750); AtPGAP1/HLD1 (At3g27325); SETH1 (At2g34980); SETH2 (At3g45100); UBQ10 (At4g05320)*.

## Supplementary Information

Materials and Methods

Figs. S1 to S14

Tables. S1 to S5

References for Materials and Methods (refs ^44-55^)

## Supplementary Information for

### Materials and Methods

#### Plant materials and growth conditions

Plants were grown as described in (22). Briefly, chlorine gas sterilized seeds were sown out on LRC2 plates (2.15g.L^-1^ MS basal salts Duchefa Biochemie, 0.1g.L^-1^ MES, pH adjusted to 5.7 with KOH, 1.0% Plant Tissue Culture Agar NEOGEN), and stored in cold room (4 °C) for three days before being moved to a growth chamber for vertical growth with continuous light emitted by white fluorescent lamps (intensity 120 µmol.m^2^.s^-1^), at 22 °C. Seedlings were transferred to Jiffy pots in soil and grown under glasshouse conditions under a 16h light/8h dark regime at 22 _o_C. Seeds were collected when the plants were completely dry and kept at room temperature (RT) or 4 °C for long-term storage.

#### EMS mutagenesis and mutants screen

T2 seeds of *At*-SI lines were collected and sown out on LRC2 plates containing BASTA (10 µg.ml^-1^). Single insertion transgenic *At*-SI lines were obtained by selecting lines showing ∼75% of BASTA resistance. Homozygous lines were obtained by selecting lines whose T3 seeds showed 100% BASTA resistance. As only ∼50 seeds could be harvested from an *At*-SI plant, selected *At*-SI lines were propagated for two more generations to collect enough seeds for EMS mutagenesis. EMS mutagenesis was carried out as described in (44). M1 *At*-SI seeds after EMS mutagenesis were sown in Jiffy pots (∼3 seeds/pot) and allowed to germinate and grow in a greenhouse. The first screen was carried out ∼10 days after flowering. Mutants showing longer siliques were selected. For those M1 plants which did not show a desired phenotype, the primary inflorescences were cut back and a second screen was carried out ∼2 weeks later. In total, ∼50,000 M1 plants were screened, from which 40 mutants were identified. After eliminating those mutants with transgene mutations, altered PrpS_1_-GFP expression levels, or pseudo-fertilizations, 12 mutants were left. These mutants were named *highlanders* (*hlds*).

To eliminate the effect of the mosaic nature of the M1 genetic background, M1 *hld* mutants (male) were backcrossed with the parent *At*-SI line (female) to obtain the backcross1 (BC1) generation of *hld* mutants. To reduce the number of background single-nucleotide polymorphisms (SNPs) caused by EMS mutagenesis, BC1 *hld* mutants were backcrossed again with the parent *At*-SI line to obtain the BC2 generation (Fig S1).

#### WGS, backcrosses and SNP analysis to identify the causal gene for the *hld1* mutants

For each of the *hld* mutant lines (*hld1-3/5/7/19/24/25*), leaf disc samples from 50 self-compatible BC2 plants were pooled together, followed by DNA extraction using a CTAB-based protocol (45), and whole genome re-sequencing (WGS) using Illumina next generation sequencing (NGS) platform. The *At*-SI line was also sequenced as the background control. SNPs were identified using SHOREmap (21). As the *hlds* were gametophytic mutations, after backcrossing with the unmutagenised parent, any unlinked SNP generated by the EMS mutagenesis segregated 1:3 in the pool of mutant individuals of the F1 population of the second backcross (BC2), whereas the causative SNP segregated 1:1 (46). Thus, calculating the SNP ratio in the BC2 population allowed us to identify the causal mutation region.

#### Cloning, transgenic and T-DNA lines

All the expression vectors were generated using either Greengate cloning (47), or Gibson assembly (New England BioLabs). High-fidelity Phusion DNA polymerase (New England BioLabs) was used for all the DNA fragment amplification. All the clones were verified through Sanger sequencing.

The expression clone pFASTGreen-pHLD1::mCherry-cHLD1 and pFASTGreen-pHLD1::mCherry-cHLD1(S218A) was generated using Greengate cloning, during which entry clones pGG-A-pHLD1-B, pGG-B-mCherry-C, pGG-C-cHLD1-D or pGG-cHLD1(S218A)-D, pGG-D-linker-E, pGG-E-G7T-F, and pGG-F-linker-G were cloned into Greengate destination vectors pFAST-GK-AG (23). The Greengate entry clone pGG-B-mCherry-C, pGG-D-linker-E, pGG-E-G7T-F, and pGG-F-linker-G were obtained from PSB Plasmid Vector Collection (https://gatewayvectors.vib.be/collection). The Greengate entry clone pGG-A-pHLD1-B was obtained through Gibson assembly. The 2755 bps upstream of At3g27325.2 translational start site was amplified as the HLD1 promoter (pHLD1) sequence. As there is a BsaI restriction site within the pHLD1 sequence, two pHLD1 fragments were separately amplified using primer sets 665/666 and 667/668, respectively, with Col-0 Arabidopsis gDNA as the template to mutate the BsaI site, and assembled into BsaI linearized pGGA000 entry vector backbone using Gibson assembly. To make the Greengate entry clone pGG-C-cHLD1-D, HLD1 cDNA (cHLD1) was amplified and cloned into pJET1.2 using CloneJET PCR Cloning Kit (ThermoFisher). There are three different HLD1 splice variants according to TAIR (https://www.arabidopsis.org) annotation. The coding region of HLD1.1 was amplified in this experiment. As there are 3 BsaI restriction sites within the coding region of cHLD1.1, cHLD1.1 was amplified through a two-step protocol. Four cHLD1.1 fragments were amplified using primer sets 732/675, 676/677, 678/679, and 680/683 with Arabidopsis seedling cDNA as the template. Mutations in these primers were designed so that the encoded protein was not changed. These four DNA fragments were assembled into cHLD1.1 using overlap PCR. The resulting PCR products were cloned into pJET plasmid vector backbone to obtain the Greengate entry clone pGG-C-cHLD1-D. To make the Greengate entry clone pGG-C-cHLD1(S218A)-D, two cHLD1(S218A) fragments were amplified using primer sets 732/834, 833/683 with pGG-C-cHLD1-D as the template. Mutations in the primers were designed so that the Serine 218 was changed to Alanine. These two PCR fragments were assembled into cHLD1(S218A) using overlap PCR, after which the resulting PCR products were cloned into pJET plasmid vector backbone to obtain the Greengate entry clone pGG-C-cHLD1(S218A)-D. Detailed primer information can be found in Supplemental Table S5. The expression vectors were transformed into GV3101 Agrobacterium tumefaciens competent cells. The floral-dipping method was adopted to stably transform homozygous *At-SI/hld1-7* Arabidopsis plants to obtain At-SI/hld1/cHLD1 and At-SI/hld1/cHLD1(S218A) lines.

To generate new HLD1 mutant alleles, 4 gRNAs (Table S5) were designed using CRISPOR and cloned into BbsI linearized Greengate entry vectors pGG-A-AtU6-26-B, pGG-B-AtU6-26-C, pGG-C-AtU6-26-D, and pGG-D-AtU6-26-E respectively via Gibson assembly. The resulting Greengate entry modules were assembled into pFASTGK-pUbi-Cas9-AG together with pGG-E-linker-G to generate expression vector pFASTGK-CRISPR-HLD1, which was transformed into *At-PrpS*_*1*_ background line via Agrobacterium-mediated floral-dipping. *Hld1-c1* and *hld1-c2* alleles were screened from T2 Cas9-free seedlings and used for further experiments.

Seeds of T-DNA lines, *seth1-2* (SAIL_674_B03) and *seth2* (SALK_039599) were obtained from The Nottingham Arabidopsis Stock Center (NASC). Primers used for genotyping these two T-DNA mutants can be found in Table S5. Genotyping of plants were carried out using Phire Plant Direct PCR Kit (Thermo Scientific) according to the manufactural instructions.

#### Triton X-114 assay, and PI-PLC assay

Three-day-old seedlings were collected into a 2 mL tube with 3 steel beads (3 mm) and frozen in liquid N_2_. Frozen samples were homogenized using a grinding mill (Retsch MM 400) at 20 Hz for 3 × 40 s. Samples were put back to liquid N_2_ after each grinding. For Triton X-114 assays, 2% Triton X-114 extraction buffer [100mM Tris-HCl pH=7.5, 150mM NaCl, 2mM EDTA, 10% glycerol, 2% Triton X-114 (Sigma-Aldrich), 1x cOmplete protease inhibitor (Roche)] was added to the homogenized samples. Triton X-114 was pre-processed as described in (49) before being added to extraction buffer. Samples were centrifuged at 21,000g at 4 °C, and the supernatant was kept for further experiments. Protein concentrations were determined using Bradford (Bio-Rad) assay by diluting the supernatant 10 times. To separate the detergent and aqueous phase, samples were incubated at 37 °C for 10 mins followed by centrifugation at 21,000g for 10 mins (50). The aqueous upper phase was carefully moved into a new tube. Proteins of both the detergent and aqueous phase were precipitated using MeOH/CHCl_3_ (51) and dissolved in 1x loading buffer.

PI-PLC assays were modified from protocols published by (52, 53). In brief, membrane protein extraction buffer [100 mM Tris-HCl pH 7.5, 25% (w/w) sucrose, 5% (v/v) glycerol, 5 mM EDTA, 5 mM KCl, 2x cOmplete protease inhibitor (Roche)] was added to the homogenized samples. Samples were centrifuged at 600g for 3 mins and the supernatants were kept as whole protein extracts. To obtain the membrane fraction, supernatants were diluted with equal volume of water and centrifuged at 21,000g for 2 hours at 4 °C. The resulting pellets were suspended in PI-PLC treatment solution [10 mM Tris-HCl pH 7.5, 5% (v/v) glycerol]. PI-PLC (ThermoFisher) was added to a final concentration of 2 units.mL^-1^. For mock control, 50% glycerol was added. Samples were incubated at 37 °C for 1.5 h, followed by centrifugation at 21,000g for 2 hours at 4 °C. The supernatant upper phase was carefully moved into a new tube, and precipitated using MeOH/CHCl3 (51). The pellet was dissolved in 1x loading buffer directly.

SDS-PAGE and Western blot were carried out as described in (19) with minor modifications. Proteins were separated by 4-20% Mini-PROTEAN TGX Stain-Free Precast gels (Bio-Rad). Stain-free signals are indicated as “loading” in the figures. The primary antibodies anti-GFP (Takara), and anti-Aquaporins (Agrisera) were used at 1:1000 dilution, and anti-α-Tubulin (Sigma-Aldrich) were used at 1:4000 dilution. The secondary HRP-conjugated anti-Mouse (GEHEALTH) and anti-Rabbit (GEHEALTH) antibodies were both used at a dilution of 1:5000. The Western blot signals were detected using WesternBright ECL HRP substrate (ADVANSTA).

#### Seedlings treated with PrsS proteins

PrsS protein treatment was performed as described in (22). Briefly, PrsS proteins were dialysed in 1/5 LRC2 liquid medium overnight in 4 °C before use. The concentration of PrsS proteins was determined using Bradford assay (BioRad). To examine the effect of PrsS proteins on seedling growth, 10 μl PrsS proteins (10 ng.µl^-1^) were added to the root tip of 4-day-old seedling on LRC2 plates. The plates were kept horizontally for 30 min to allow the PrsS proteins to dry before being placed back to the growth chamber vertically.

#### Arabidopsis pollen SI bioassay *in-vitro*

Arabidopsis pollen was transferred onto the surface of an 8-well Chambered Coverglass (Thermo Scientific Nunc Lab-Tek) by using tweezers to hold a stage 13-14 flower inverted of the coverglass and brushing the flower over the coverglass gently. Without hydrating the pollen, *A. thaliana* pollen germination medium [AtGM; 15% (w/v) sucrose, 0.01% (w/v) H_3_BO_3_, 5 mM KCl, 1 mM MgSO_4_, 2.5 mM CaCl_2_, 2.5 mM Ca(NO_3_)_2_, and 10% PEG3350, pH=7.0 adjusted using KOH; modified from (54) containing 10 ng.µl^-1^ PrsS proteins (SI treatment) or not (Mock control)] was added. For each sample, 200 µl medium was used. Samples were incubated at RT for 6h before FDA/PI co-staining and fluorescent microscopy examination. FDA (2 μg.ml^-1^) and PI (5 μg.ml^-1^) were added to the sample right before microscopy examination, during which the number of FDA positive (alive) and PI positive (dead) pollen were counted. For each sample, 100 - 200 pollen were counted. When the PLC inhibitors (U73122, ET-18-OCH3, C48/80) were needed, the pollen were pretreated with the PLC inhibitors for 1h before the SI was induced. U73122 (Sigma-Aldrich) was dissolved in chloroform. ET-18-OCH3 (Sigma-Aldrich) was dissolved in DMSO, and C48/80 (Sigma-Aldrich) was dissolved in water. For mock treatment, equal amount of solvent was used.

#### PGAP1 Phylogeny analysis

PGAP1 domain containing proteins were downloaded from Pfam (http://pfam.xfam.org/family/PGAP1). In total, 631 proteins from species across three eukaryotic kingdoms were downloaded (Table S6). Multiple sequence alignment was performed using MAFFT (version 7.187). IQ-TREE (version 1.7-beta7) was used for the maximum-likelihood tree inference with 1000 bootstrap replicates (under the model of ‘JTT+R’, -alrt 1000 -bb 1000).

#### Imaging, image analysis and figure preparation

Imaging of the PI stained root was performed using a Zeiss LSM710 microscope using a PlanApochromat 20x objective (numerical aperture 0.8). Seedling samples were mounted with 1/5 LRC2 medium containing 5 μg.ml^-1^ PI. PI was excited with 561 nm and fluorescence emission between 580 nm and 700 nm were collected.

For the *in-vitro* pollen SI bioassay, pollen was co-stained using FDA (2 μg.ml^-1^) and PI (5 μg.ml^-1^) and visualized using a Leica SP8 confocal laser scanning system with Fluostar VISIR 25x/0.95 water objective and HyD detector. FDA was excited with 488 nm and fluorescence emissions between 500 nm and 550 nm were collected. PI was excited with 561 nm and fluorescence emission between 580 nm and 700 nm were collected.

GFP-SKU5 were visualized using a Leica SP8 confocal laser scanning system with HCPL APO CS2 40x/1.10 (water) objective and HyD detector. Samples were excited with 488 nm, and fluorescence emissions between 500 nm and 550 nm were collected.

All the images were processed and analyzed using Fiji (https://fiji.sc/) (55).

#### Statistical analysis

Statistical analysis was performed using GraphPad Prism 8.0 for Windows (www.graphpad.com).

**Supplemental Figure S1.**
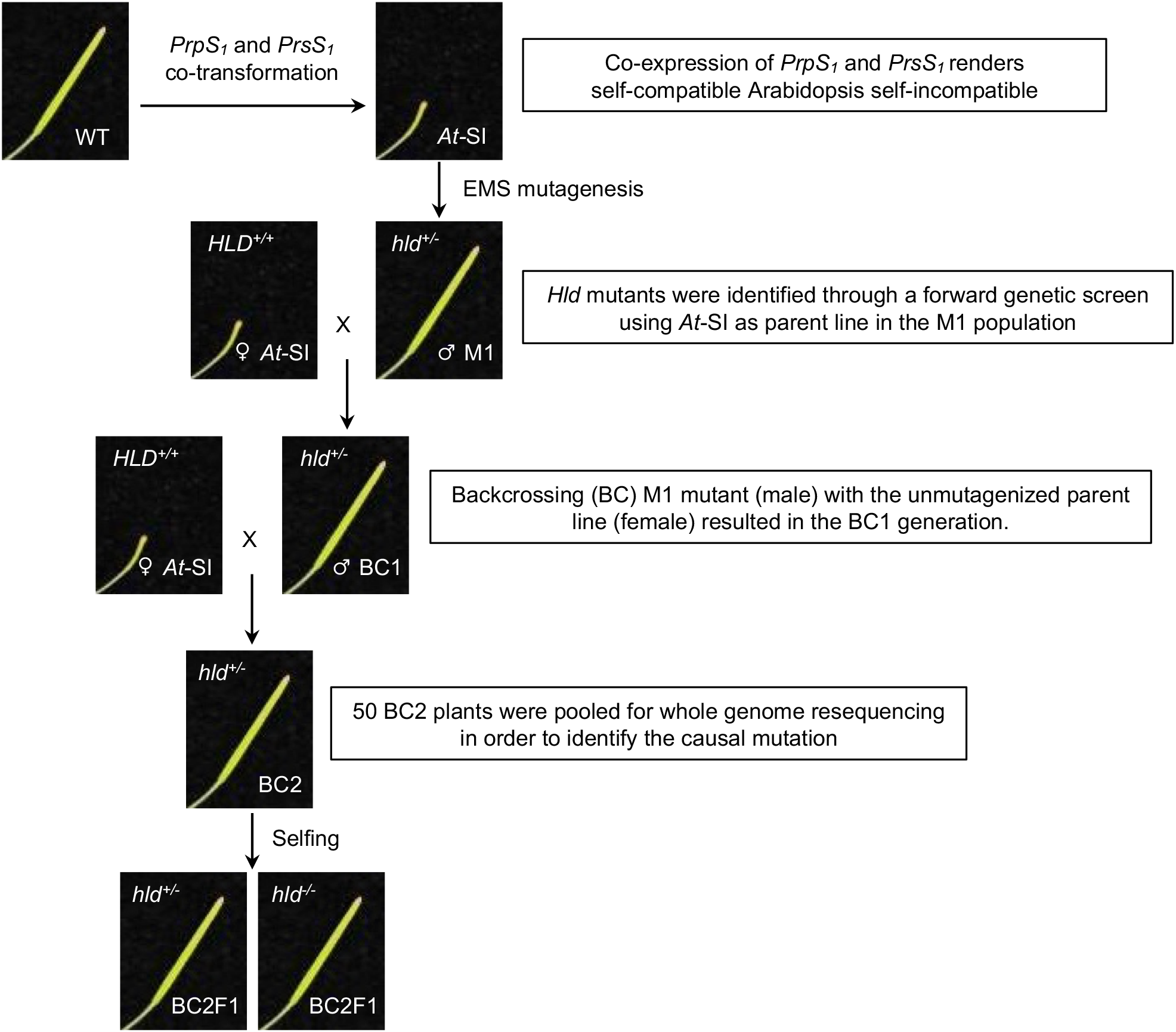
Experimental scheme. Co-expression of *PrpS*_*1*_ and *PrsS*_*1*_ in the Col-0 background renders self-compatible Arabidopsis self-incompatible. The resulting SI Arabidopsis was used as the parent line in an EMS-based forward genetic screen looking for SC mutants. The mutants were named *highlanders* (*hlds*). *Hld* mutants identified in the M1 population were back-crossed with parent *At*-SI line to obtain the backcross 1 (BC1) generation. To reduce the number of background non-causal SNPs, BC1 plants were back-crossed again with the parent *At*-SI line to obtain the BC2 generation. Fifty BC2 plants were pooled for WGS to identify the causal mutation. Seeds collected from BC2 selfing (BC2F1) were used for the identification of homozygous *hld* mutants.

**Supplemental Figure S2.**
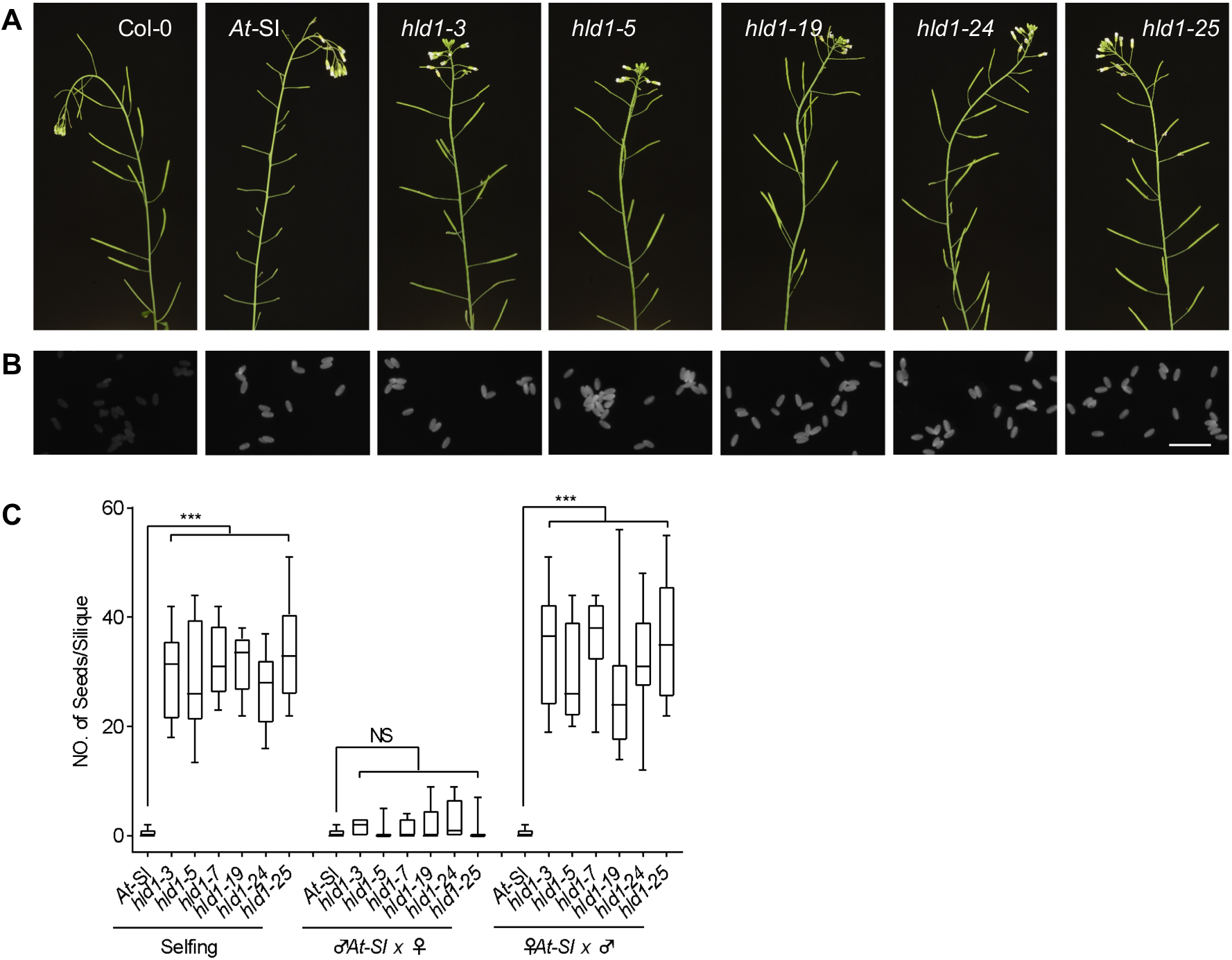
*hld1* mutants are defective in SI. (A) ***hld1* was identified from an EMS mutagenesis screen**. Mutants were screened for a defective SI phenotype, with normal seed set. Representative images of mature inflorescences from Col-0, *At*-SI, and *hld1-3/5/19/24/25* mutant plants show that all these heterozygous mutants displayed no other obvious developmental phenotype in comparison with Col-0. (B) **There was no alteration of PrpS**_**1**_**-GFP signals in *hld1* mutants**. *At*-SI pollen grains showed homozygous PrpS_1_-GFP signal while only background fluorescence was observed in Col-0. No gross alteration in the PrpS_1_-GFP signals was observed in any of the *hld1* mutants compared to the parental *At*-SI line. Bar = 100 μm. (C) ***hld1* mutants have pollen defects**. *hld1* stigmas were pollinated with self-pollen (selfing) or *At-* SI pollen (♂At-*SI* x ♀) and *At-*SI stigmas pollinated with *hld1* pollen (♀At-*SI* x ♂). Silique length measurements (Fig 1B) and number of seeds per silique (C) showed that the *hld1* mutants used as the male parent had significantly longer silique lengths and higher seed-set than the *At-*SI parent line. *hld1* mutants used as the female parent pollinated with *At*-SI pollen displayed a normal SI phenotype of short silique lengths and low seed set. N=9-11. One-way ANOVA. ***: p<0.001. NS: not-significant, p>0.05.

**Supplemental Figure S3.**
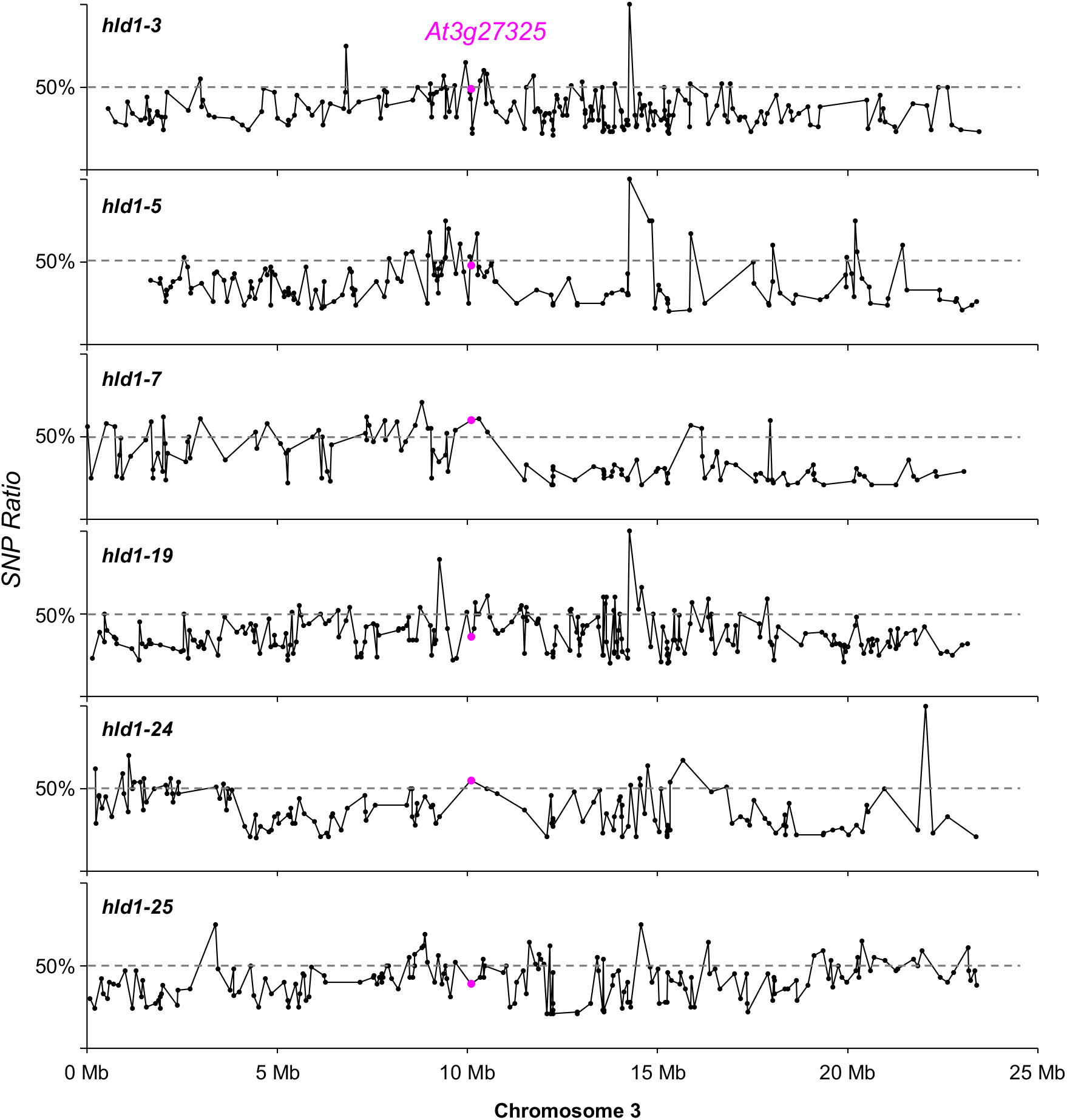
WGS identified *At3g27325* as the causal gene candidate of the *hld1* mutants. As the *hlds* were gametophytic mutations, after backcrossing with the unmutagenised parent, any unlinked SNP generated by the EMS mutagenesis segregated 1:3 in the pool of mutant individuals of the F1 population of the second backcross (BC2), whereas the causative SNP segregated 1:1. Thus, calculating the SNP ratio in the self-compatible BC2 population allowed us to identify the causal mutation region. Analysis of the SNPs in the 6 *hld1* mutants uncovered six different nonsynonymous mutations of an uncharacterized gene, *At3g27325*, in all the 6 different *hld1* mutants. This suggests that *At3g27325* is the causal gene responsible for the SC phenotype in all the 6 *hld1* mutants.

**Supplemental Figure S4.**
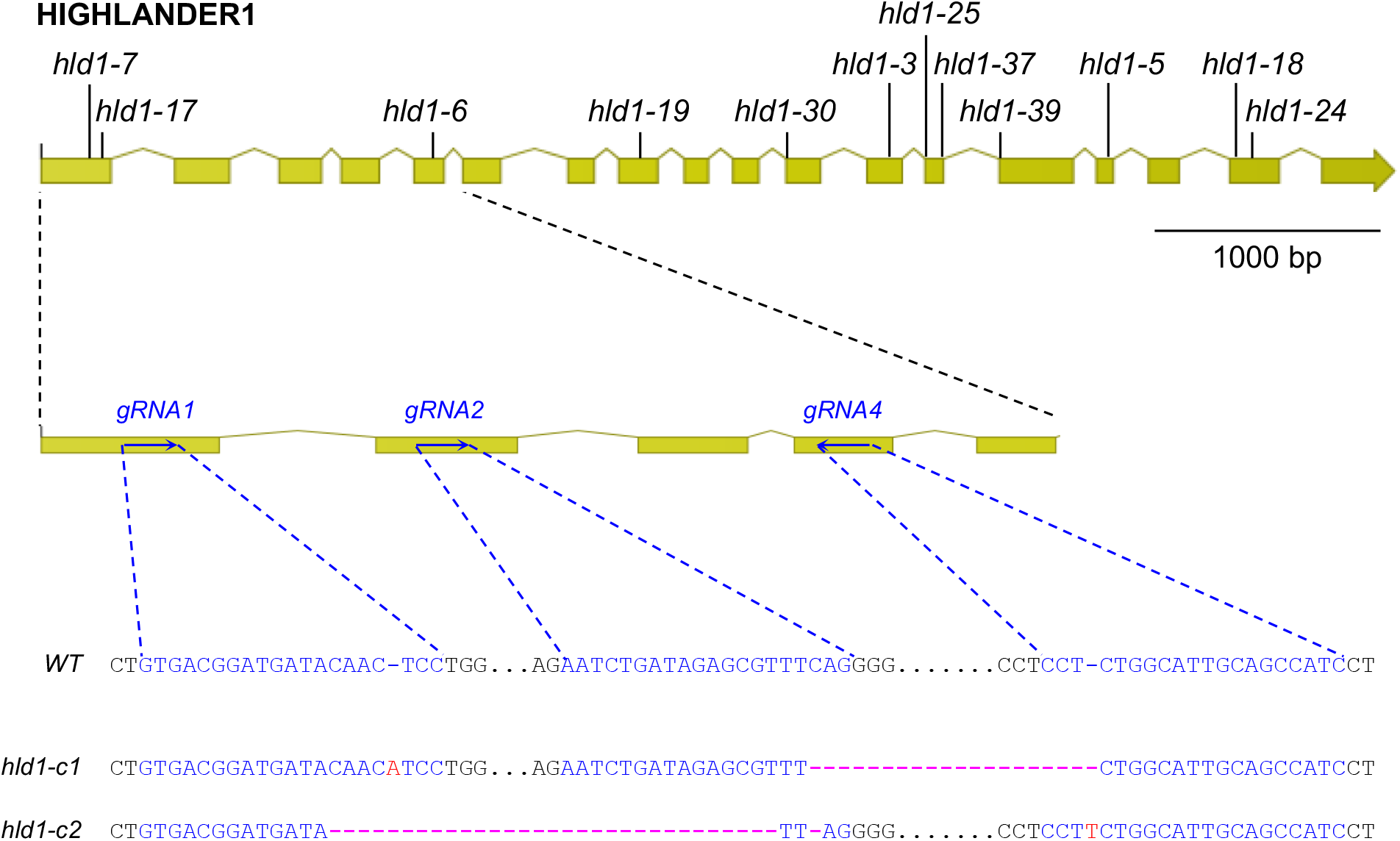
Two new *At3g27325* mutant alleles were generated using CRISPR/Cas9. *Hld1-3/5/6/7/17/18/19/24/25/30/37/39* (upper row) were identified from the EMS screen; *hld1-c1* and *hld1-c2* (bottom rows) are the CRISPR/Cas9 generated mutants. The detailed positional information of each mutant can be found in **Figure 1C** and **Supplemental Table S3**. Yellow boxes indicate the exons of *At3g27325.1*, blue text indicates gRNAs, red nucleotides indicate insertions, pink dashes indicate deletions.

**Supplementary Figure S5.**
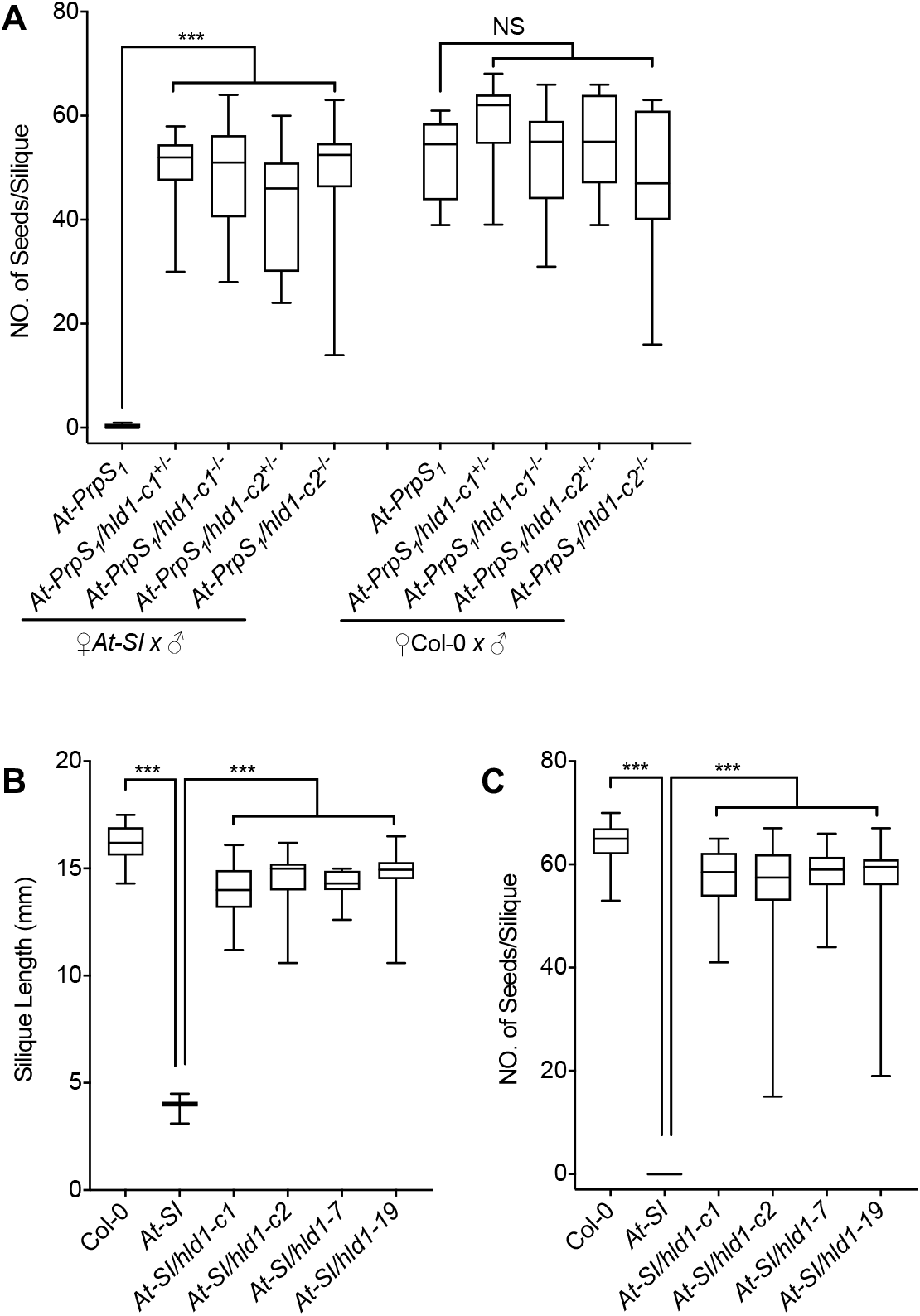
Evidence that *At3g27325* is the causal gene of the *hld1* mutants. (A-C) **Mutation of *At3g27325* phenocopies *hld1***. (A) *At-PrpS*_*1*_ pollen containing the CRISPR/Cas9-derived *At3g27325* mutant allele (*hld1-c1* or *hld1-c2*) were pollinated onto *At-*SI or Col-0 stigmas. Significant increases in the silique lengths (**Figure 1D**) and number of seeds per silique (A) were observed when *At3g27325* was mutated, in both heterozygous and homozygous mutants. N=12-16. One-way ANOVA with multiple comparisons. ***: p<0.001. NS: p>0.1772. (B-C) Introduction of the *hld1-c1* or *hld1-c2* mutant allele into the *At*-SI background resulted in SC mutants showing silique lengths (B) and seed-set levels (C) similar to the EMS mutagenesis generated *hld1* mutants. N = 30 siliques from 3 independent plants, 10 each. One-way ANOVA with multiple comparisons. ***: p<0.001.

**Supplementary Figure S6.**
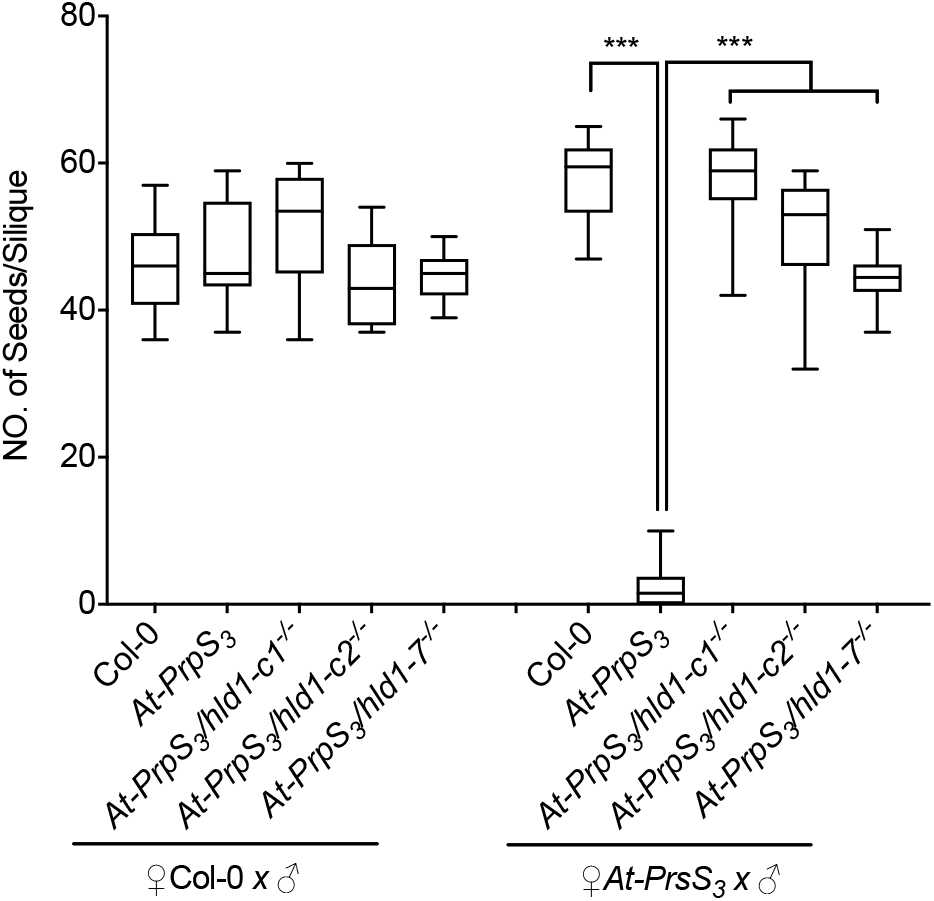
*HLD1* regulates SI in an *S*-specific manner. (A-C) *At-PrpS*_*3*_ pollen containing the *hld1* mutant allele (*hld1-c1, hld1-c2*, or *hld1-7*) were pollinated onto *At-PrsS*_*3*_ or Col-0 stigmas. Silique lengths (**Figure 2A**) and seed set were measured. Consistent with previous observations, when *Col-0* stigmas were pollinated with *Col-0*, or *At-PrpS*_*3*_, or *At-PrpS*_*3*_*/hld1* pollen, there was no significant difference observed in the silique length and number of seeds per silique (*20*). Pollinating *At-PrsS*_*3*_ stigmas with *At-PrpS*_*3*_ pollen resulted in SI, with a significant reduction in silique length and almost no seed, compared with the control pollination ♀*At-PrsS*_*3*_ x ♂*Col-0*. In contrast, pollinating *At-PrsS*_*3*_ stigmas with *At-PrpS*_*3*_*/hld1* pollen resulted in silique length and seed-set like the controls, demonstrating that mutation of the *HLD1* gene abolishes the *PrpS*_*3*_*-PrsS*_*3*_-based SI. This provides good evidence that *HLD1* acts as a SI regulator in an *S*-specific manner. N=10-20. One-way ANOVA. ***: p<0.001.

**Supplemental Figure S7.**
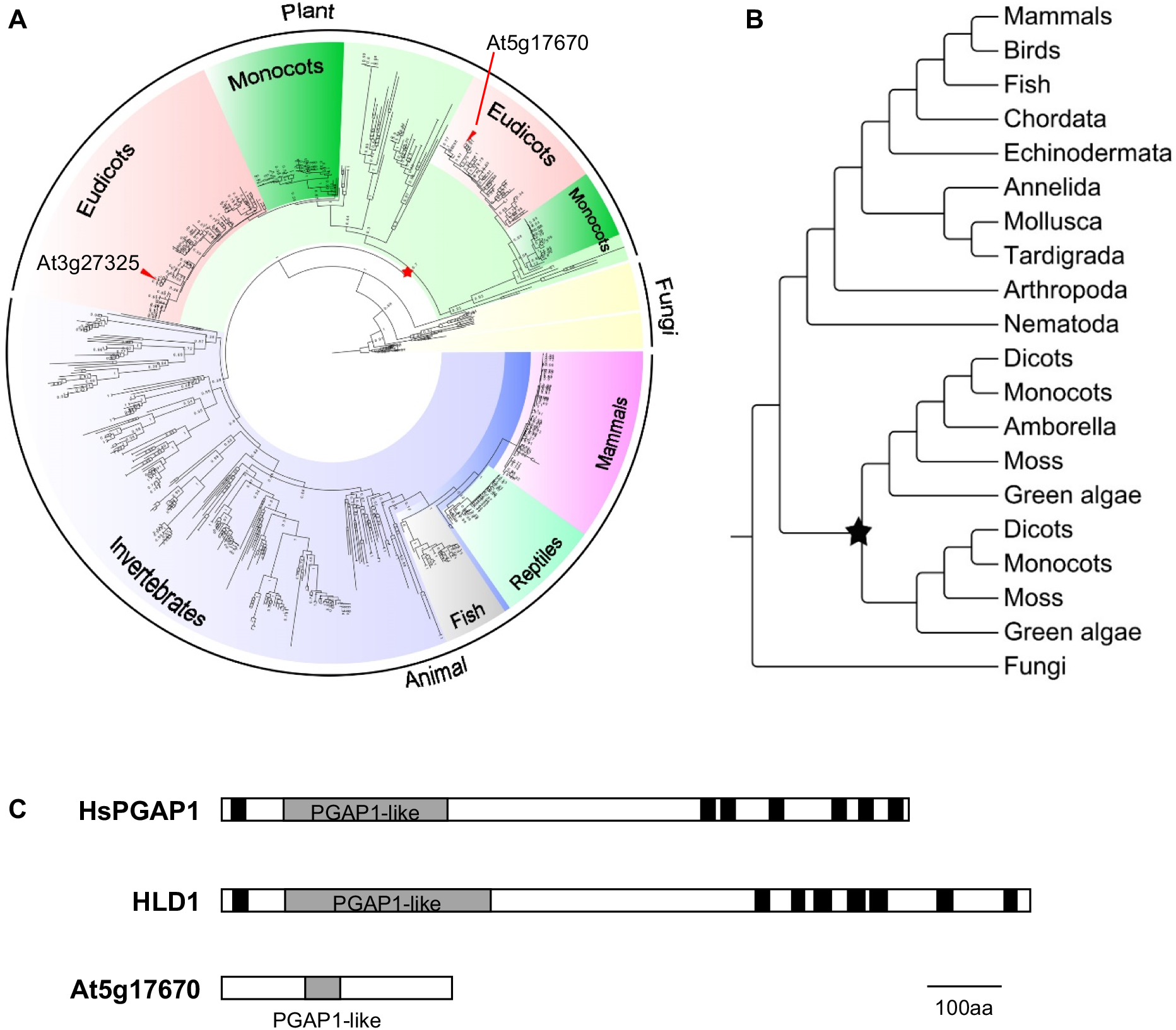
HLD1 is a HsPGAP1 orthologue. (A-B) **Phylogenetic analysis of the PGAP1-like domain-containing proteins across eukaryotic kingdoms**. We constructed a phylogenetic tree of 631 predicted PGAP1 protein homologues from > 300 eukaryotic species. Two major phylogenetic clades were identified for plants; in Arabidopsis, *HLD1* and another *PGAP1* homologue *At5g17670* were classified into different clades, indicated by red triangles. The divergence of these two clades can be traced back to an ancient whole genome duplication event occurring before the radiation of extant Viridiplantae, including green algae and the land plants. Stars indicate genome duplication events. Detailed information of the 631 PGAP1-like domain-containing proteins used in the construction of the phylogeny tree can be found in Supplemental Table S6. (C) **Cartoon of the predicted secondary structure of HsPGAP1, HLD1 and At5g17670**. Comparison of the two homologs, *HLD1* and *At5g17670* from *A. thaliana* with *Homo sapiens PGAP1*, revealed that *HLD1* has a slightly higher amino acid sequence similarity to *HsPGAP1* (23.3%) than *At5g17670* (20.1%). Importantly, *HLD1* shares a similar secondary protein structure to *HsPGAP1* while *At5g17670* does not. Grey boxes represent the PGAP1-like domain. Black boxes represent predicted transmembrane domains. This suggests that *HLD1* is likely to be the *HsPGAP1* ortholog in Arabidopsis.

**Supplemental Figure S8.**
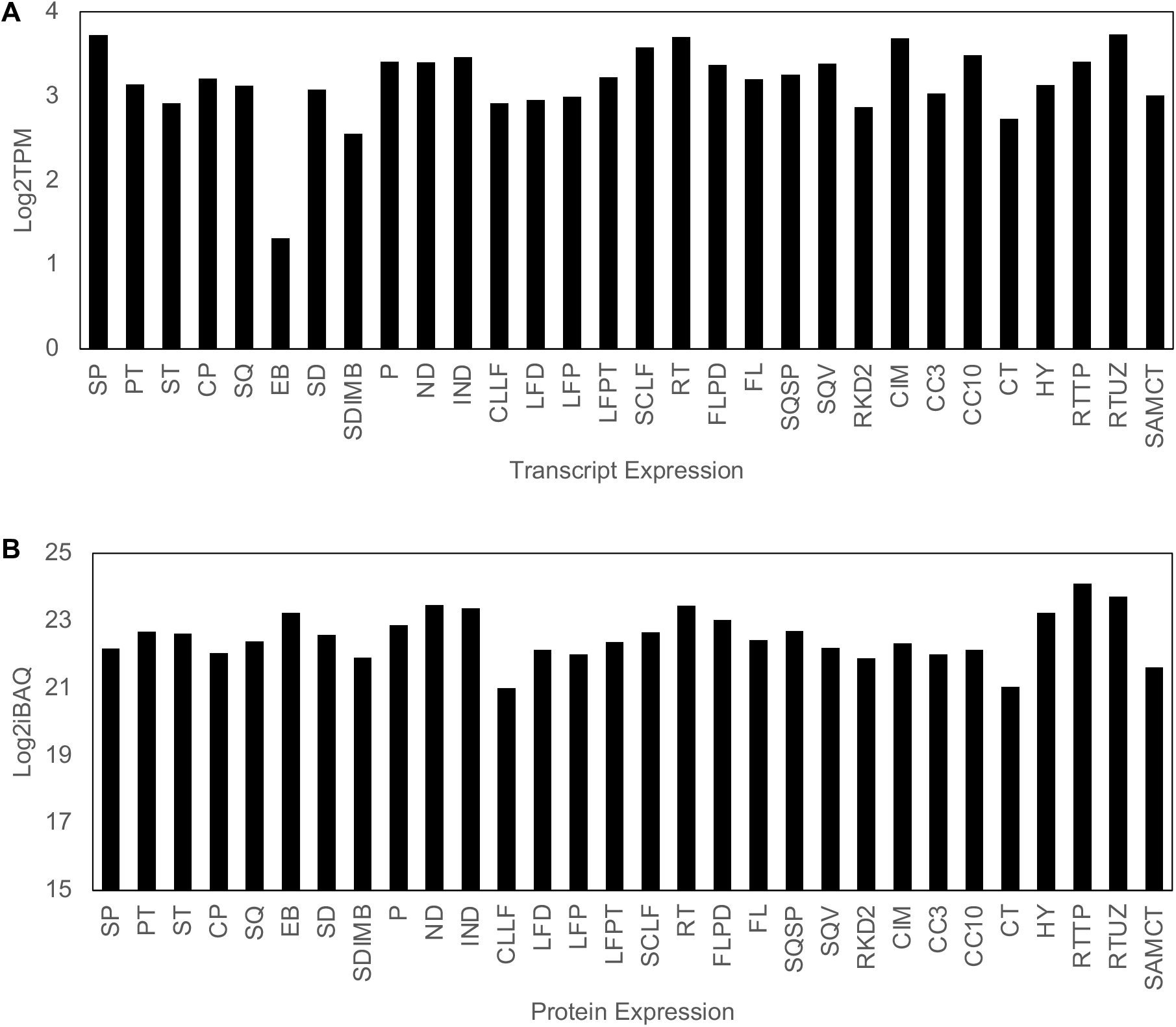
Transcript and protein expression patterns of HLD1 (At3g27325) in *Arabidopsis thaliana* tissues. The transcript (A) and protein (B) expression of the *HLD1* gene (*At3g27325*) across 30 different tissues was investigated using publicly available transcriptome and proteomics data (http://athena.proteomics.wzw.tum.de/). TPM: transcripts per million. iBAQ: intensity-based absolute quantification. Abbreviations: Sepal (SP); Petal (PT); Stamen (ST); Carpel (CP); Silique (SQ); Embryo (EB); Seed (SD); Seed imbibed (SDIMB); Pollen (P); Node (ND); Internode (IND); Cauline leaf (CLLF); Leaf distal (LFD); Leaf proximal (LFP); Leaf petiole (LFPT); Senescent leaf (SCLF); Root (RT); Flower pedicle (FLPD); Flower (FL); Silique septum (SQSP); Silique valves (SQV); Egg-cell like callus (RKD2); Callus (CIM); Cell culture early (CC3); Cell culture late (CC10); Cotyledons (CT); Hypocotyl (HY) ; Root tip (RTTP); Root upper zone (RTUZ); Shoot tip (SAMCT).

**Supplemental Figure S9.**
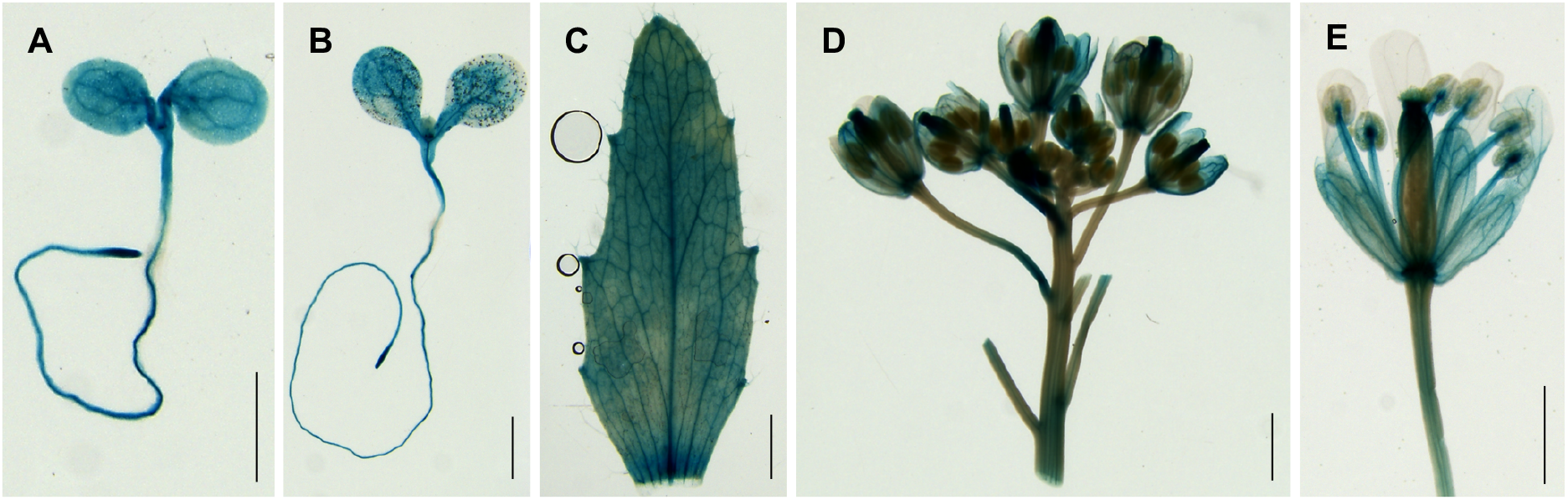
HLD1 is expressed in various plant tissues. The HLD1 expression pattern was examined using a pHLD1::GUS transgenic line. HLD1 was expressed in all the plant tissues examined. (A) 5-day-old seedling; (B) 8-day-old seedling; (C) mature leaf; (D) flower bud; (E) stage 13 flower. Representative images are shown from at least three independent lines. Bar = 1 cm.

**Supplemental Figure S10.**
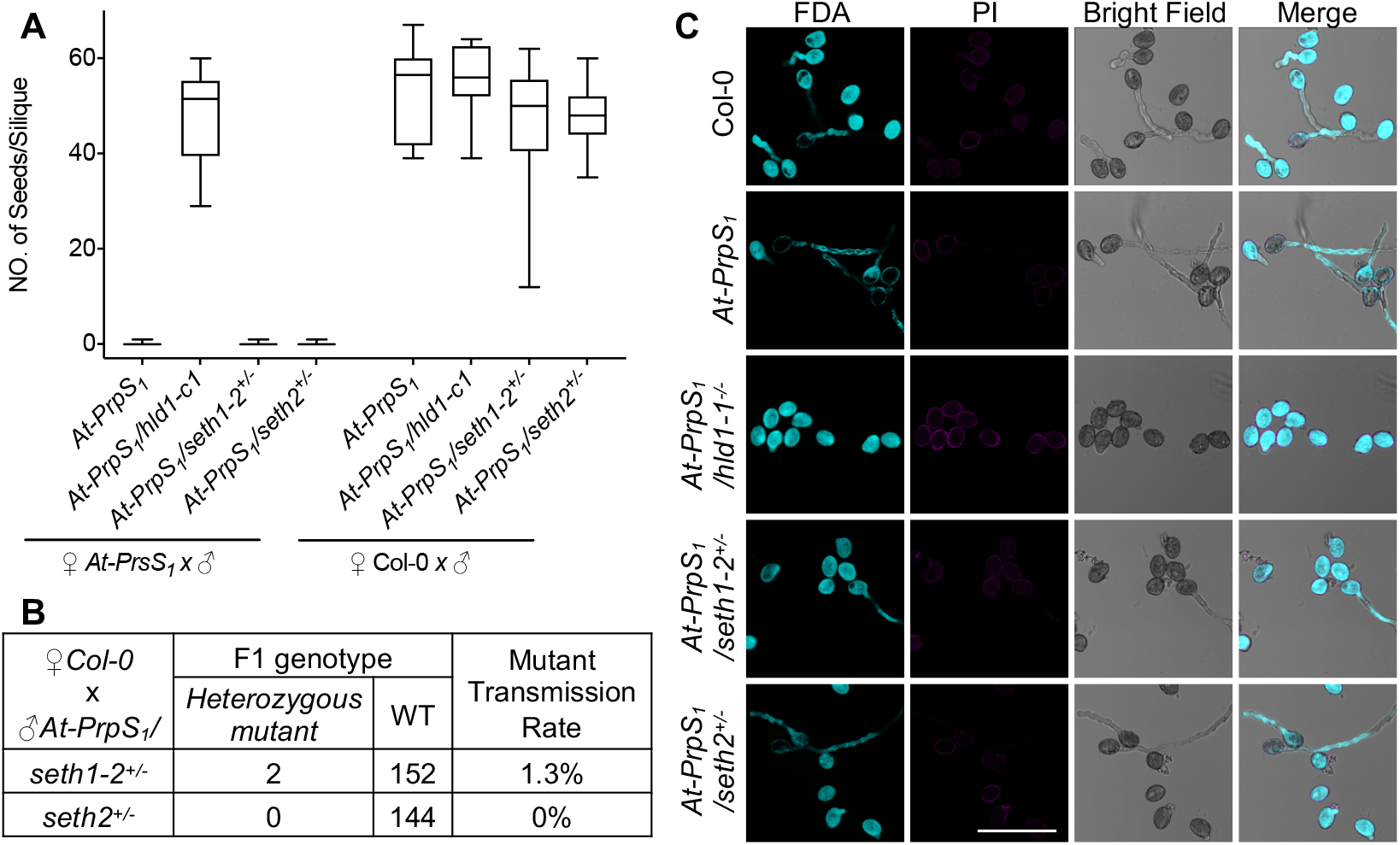
Evidence for involvement of the GPI-anchoring pathway in the regulation of SI-induced death of pollen. (A-B) **Mutation of GPI-anchoring pathway genes does not phenocopy the *hld1* mutant phenotype**. GPI-anchoring pathway mutants, *seth1-2* and *seth2*, were introduced into *At-PrpS*_*1*_ background, and the pollen were pollinated onto *At-PrsS*_*1*_ or Col-0 stigmas. Pollinating *At-PrsS*_*1*_ stigmas with *At-PrpS*_*1*_*/seth1*^*+/-*^ or *At-PrpS*_*1*_*/seth2*^*+/-*^ pollen resulted in short siliques (Fig 4A) and no seedset (A). N=13-17. (B) Genotyping the F1 seedlings of ♀*Col-0* x ♂*At-PrpS*_*1*_*/seth1*^*+/-*^ and ♀*Col-0* x ♂*At-PrpS*_*1*_*/seth2*^*+/-*^ revealed that mutation of *seth1* or *seth2* abolished pollen fertilization completely. (C) **Mutation of GPI anchor synthesis did not affect pollen viability**. The *seth1-2* and *seth2* mutations were introduced into *At-PrpS*_*1*_ background, and the pollen was mock treated with *At*-GM (C) or PrsS_1_ proteins (Fig 4B). The samples were co-stained with FDA and PI 6h after treatment. All the pollen samples showed >97% viability when treated with *At*-GM buffer. When treated with PrsS_1_ proteins (Fig 4B), only minimal (∼3%) pollen death was observed for *Col-0* or *At-PrpS*_*1*_*/hld1* pollen, which was comparable to that of *At*-GM buffer control. Bar = 100 μm. Three independent experiments were carried out. In each experiment, 100-200 pollen grains were counted for each of the samples.

**Supplemental Figure S11.**
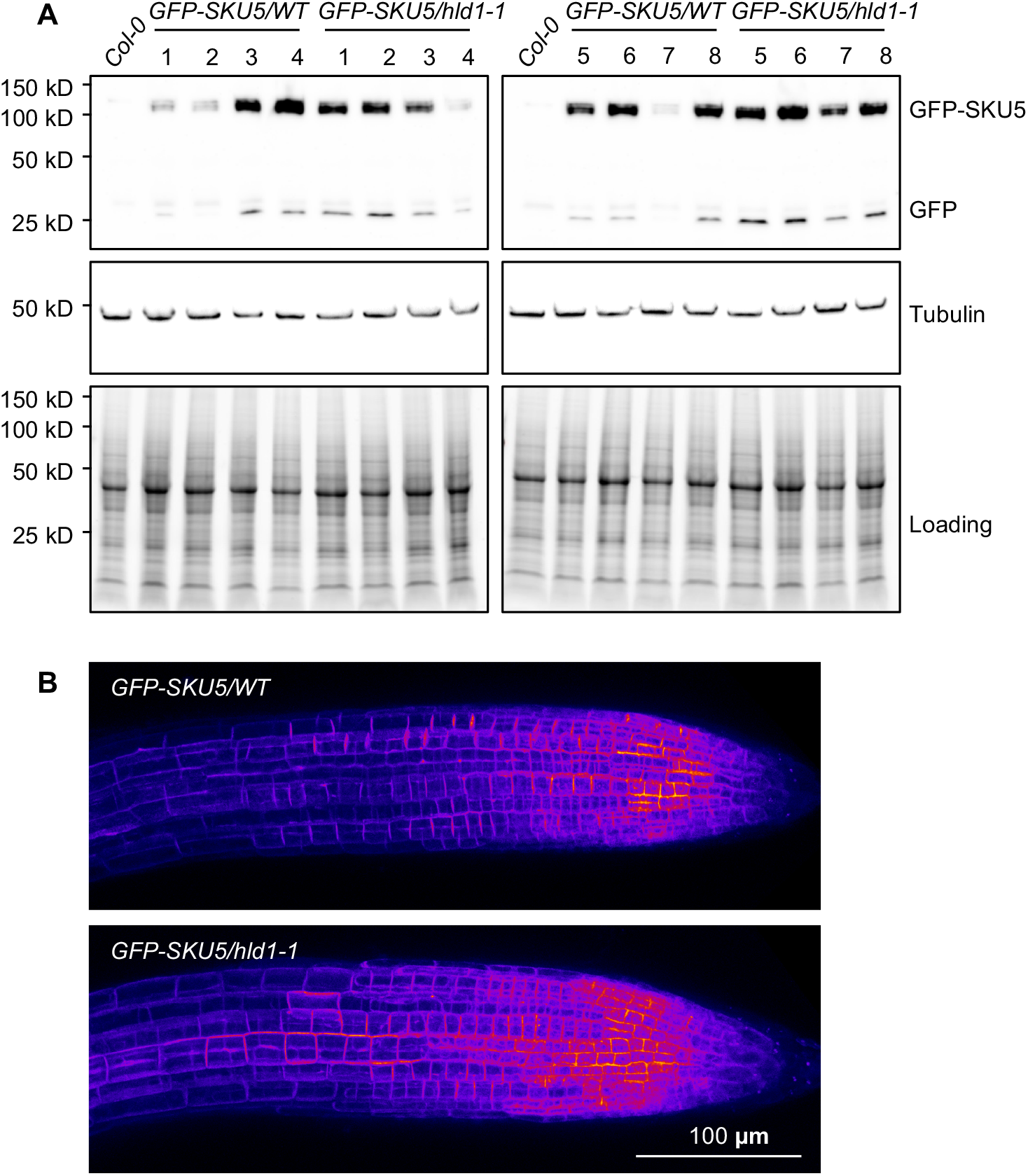
Mutation of HLD1 does not affect the GFP-SKU5 protein expression level and localization. (A) We examined the protein expression level of GFP-SKU5 using Western blot (A) in F3 seedlings derived from 16 F2 double homozygous siblings of the cross ♀hld1-1 x ♂GFP-SKU5. Tubulin was employed as the internal control. Quantification of the Western blot signals (Figure 5A) revealed that the GFP-SKU5 protein expression level was not altered by the HLD1 mutation (p=0.7995). (B) Representative confocal images (maximum Z projections) of GFP-SKU5 signals of 4-day-old *GFP-SKU5/WT* or *GFP-SKU5/hld1-1* seedlings. Single Z slice in Fig 5B. Confocal examination showed that the HLD1 mutation had no obvious effect on the GFP-SKU5 signal distribution.

**Supplemental Figure S12.**
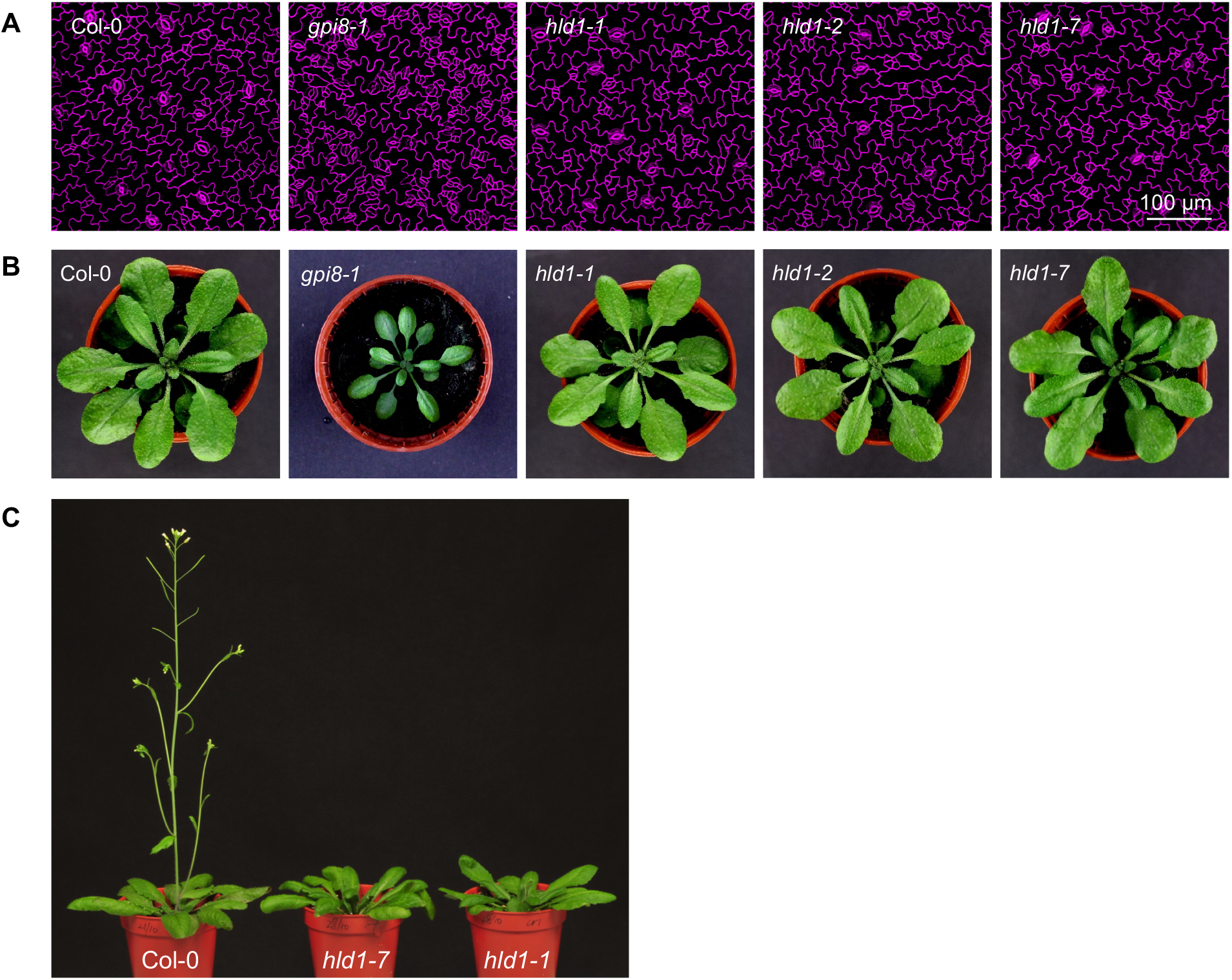
Mutation of HLD1 has no major obvious effect on plant development except for a delay in flowering time. (A-B) **Mutation of HLD1 has no major obvious effect on plant development**. The *gpi8-1* mutant was established to have reduced accumulation of GPI-APs; this affects both stomata formation, and plant growth (37). This *gpi8-1* was employed as a negative control here. (A) The *gpi8-1* mutation resulted in the formation of stomata clusters, while this was not observed in Col-0, nor the *hld1* mutants. (B) The *gpi8-1* mutation resulted in a significant delay of plant development, while this was not observed in Col-0, nor the *hld1* mutants. (C) **Mutation of HLD1 resulted in a delay in flowering time of ∼1 week**. Col-0 and homozygous *hld1* seeds were sown and grown under exactly the same conditions. Thirty-four-day-old Col-0 plants displayed extended inflorescences with flowers, while plants of the same age with the homozygous *hld1* mutation were just starting to flower. Flowering of *hld1* mutants was delayed for ∼7 days. This was observed in 3 independent experiments. Eight plants of each genotype were grown in each experiment.

**Supplemental Figure S13.**
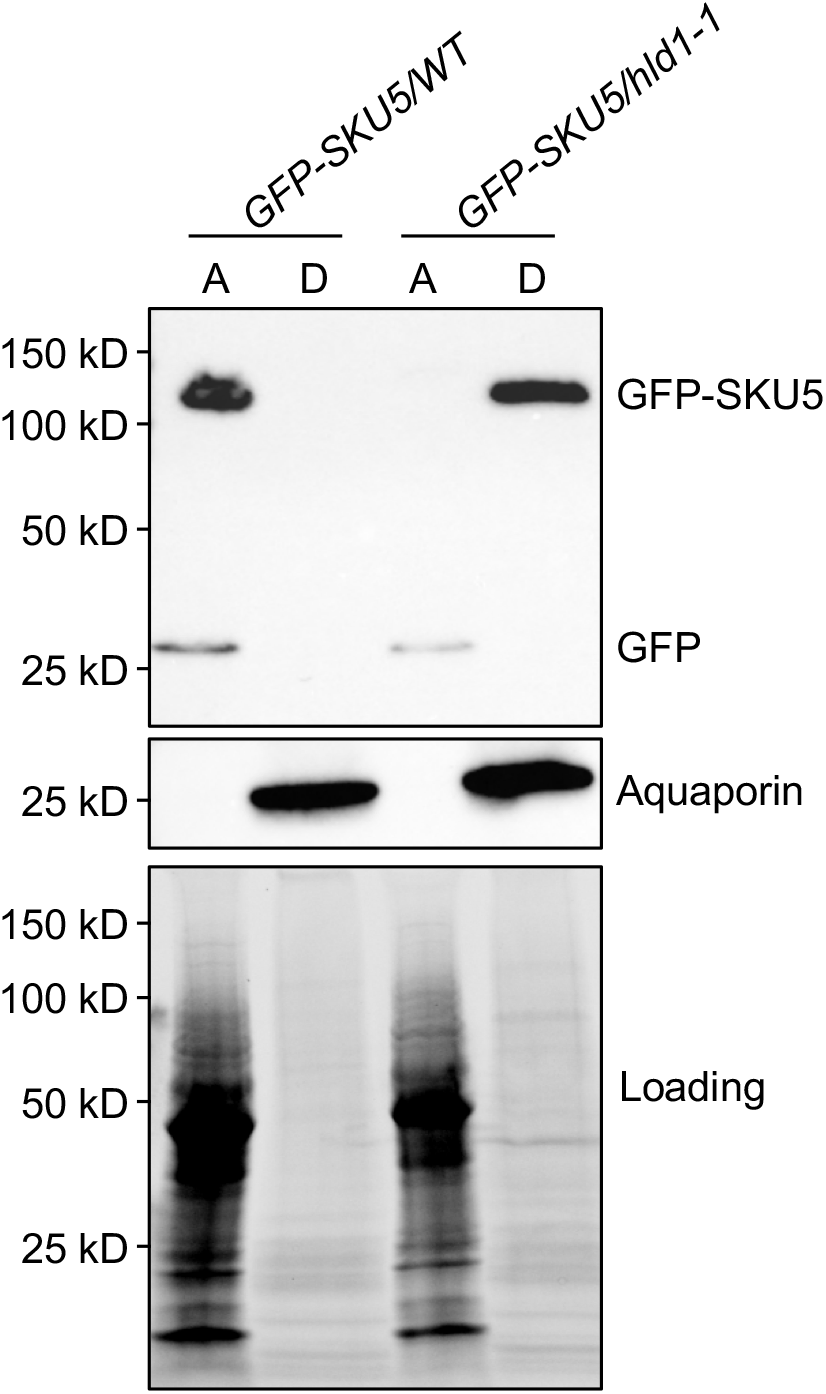
In the HLD1 mutant, an increase in hydrophobicity of GFP-SKU5 is observed. GFP-SKU5/WT and GFP-SKU5/hld1 proteins were extracted using 2% Triton X-114 extraction buffer, partitioned into aqueous phase and detergent phases, and subjected to Western blot analysis. The HLD1 mutation resulted in a shift of GFP-SKU5 from aqueous phase to detergent phase, revealing that mutation of HLD1 resulted in an increase of GFP-SKU5 hydrophobicity. A: aqueous phase; D: detergent phase.

**Supplemental Figure S14.**
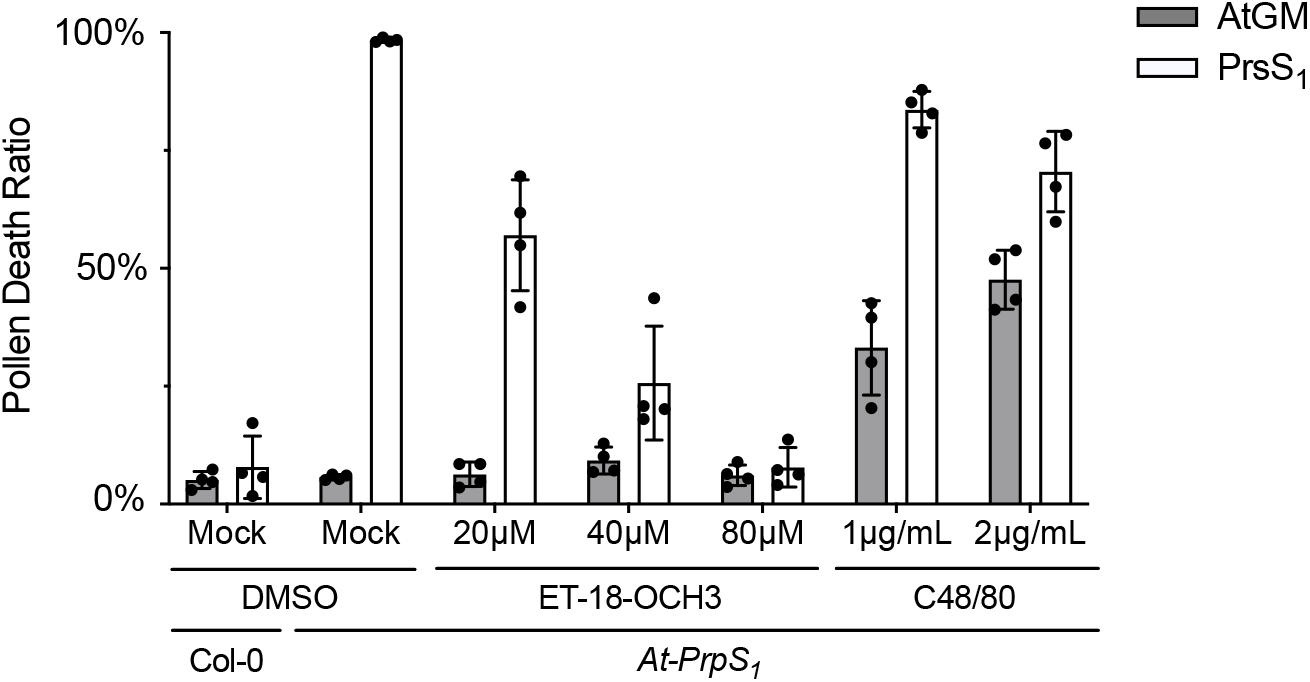
Treatment of PLC inhibitors, ET-18-OCH3 and C48/80, inhibits SI-induced pollen death. Col-0 and *At-PrpS*_*1*_ pollen grains were pretreated with different concentrations of PLC inhibitor ET-18-OCH3, C48/80, or solvent (Mock) before subject to SI induction (PrsS_1_) or control treatment (AtGM). The samples were co-stained with FDA and PI 6h after treatment. The PI positive ratios were quantified in. Four independent experiments were carried out. In each experiment, 100-200 pollen grains were counted for each of the sample. See Fig 5D for the results of another PLC inhibitor U73122.

**Supplemental Table S1.**
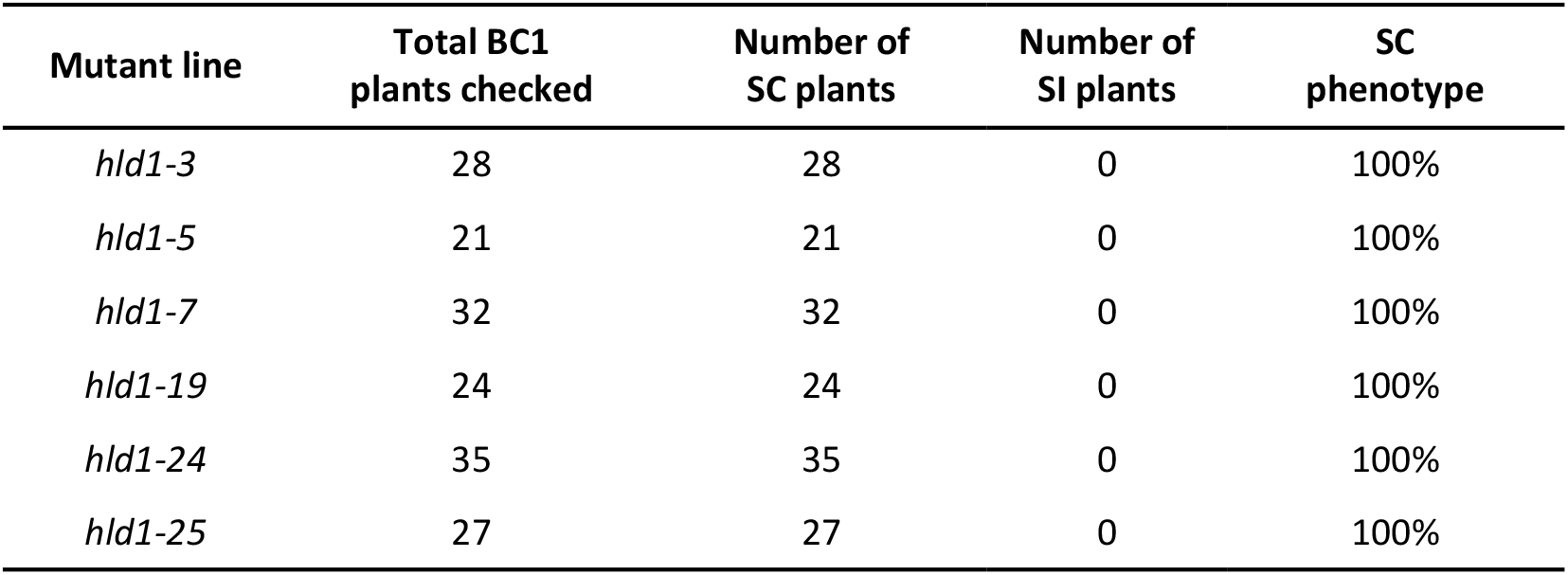
Screening of the BC1 population. After back-crossing (BC) using M1 mutant pollen with unmutagenized *At*-SI stigma, the resulting BC1 seeds were sown. The siliques of the BC1 plants were scored for either a self-compatible (SC) phenotype, showing long siliques and normal seed set, or a self-incompatible (SI) phenotype with short siliques and ∼no seed set. Examination of all 167 BC1 plants revealed that none of them showed the SI phenotype, with 100% of the BC1 plants having a SC phenotype, indicating genuine revertant mutants were identified. This fits our prediction that all the BC1 plants should be self-compatible, as in the back-cross using M1 mutant pollen on an unmutagenized *At*-SI stigma, WT pollen would undergo SI-PCD triggered by a normal PrpS_1_-PrsS_1_ interaction, so only pollen carrying a *hld1* mutation (defective in SI) would be able to fertilize successfully and produce seeds. As the *hld1* SC phenotype existed in all the heterozygous BC1 *hld1* mutant lines examined (n= 167), this demonstrates that the hld1 mutants are gametophytic.

**Supplemental Table S2.** Segregation analysis of the BC2F1 population. Back-cross 2 (BC2) plants were allowed to self, generating a BC2 F1 population. The *At*-SI parent line we selected for the EMS mutagenesis screen was leaky, and not 100% SI, otherwise, the screen would have been impossible. This meant that very few self-incompatible plants could be found in the BC2 F1 population. Resultant progeny were scored for either a self-compatible (SC) phenotype, showing long siliques and normal seed set, or a self-incompatible (SI) phenotype with short siliques and ∼no seed set. We also scored the genotype (*hld1*^+/+^, *hld1*^+/-^, *or hld1*^-/-^) of the resulting BC2 F1 population using Sanger sequencing. This identified 100% linkage between the *At3g27325* mutation and the self-fertile (SC) phenotype. This indicates that *At3g27325* is highly likely to be the *hld1* causal gene.

| mutant | phenotype | genotype | number of plants |
| --- | --- | --- | --- |
| *hld1-3* | SC | *At3g27325*<sup>+/-</sup> or *At3g27325*<sup>-/-</sup> | 7 |
|  | SI | Not found | 0 |
| *hld1-5* | SC | *At3g27325*<sup>+/-</sup> or *At3g27325*<sup>-/-</sup> | 7 |
|  | SI | Not found | 0 |
| *hld1-7* | SC | *At3g27325*<sup>+/-</sup> or *At3g27325*<sup>-/-</sup> | 197 |
|  | SI | *At3g27325*<sup>+/+</sup> | 4 |
| *hld1-19* | SC | *At3g27325*<sup>+/-</sup> or *At3g27325*<sup>-/-</sup> | 54 |
|  | SI | *At3g27325*<sup>+/+</sup> | 1 |
| *hld1-24* | SC | *At3g27325*<sup>+/-</sup> or *At3g27325*<sup>-/-</sup> | 26 |
|  | SI | *At3g27325*<sup>+/+</sup> | 2 |
| *hld1-25* | Not checked |  |  |

**Supplemental Table S3.** Identification of 6 additional *At3g27325* mutant alleles using Sanger sequencing. Genotyping of the *At3g27325* DNA region of the remaining *hld1* mutants using Sanger sequencing identified 6 additional *At3g27325* mutant alleles. It is worth to notice that for *hld1-17*, instead of a point mutation, a thymine insertion was identified, which is rarely observed in EMS mutagenesis screens. As our screen revealed 12 independent mutant alleles of *HLD1*, but no other mutants, it is likely that other genes involved in SI signaling act redundantly or are key to pollen fertilization, thus cannot be identified by forward genetic screens.

| mutant | Mutation at the DNA level | Mutation in protein level |
| --- | --- | --- |
| *hld1-6* | G1700A | W335* |
| *hld1-17* | C234CT | K80* |
| *hld1-18* | C5181T | Q918* |
| *hld1-30* | G3223A | Splice site change |
| *hld1-37* | G3913A | Splice site change |
| *hld1-39* | G4145A | Splice site change |

**Supplemental Table S4.** HLD1 lipase activity is required for the SI response. *Hld1* mutants were transformed with *pHLD1::mCherry-cHLD1* or *pHLD1::mCherry-cHLD1(S218A)* cloned in the pFAST-Green plasmid vector backbone. The pollen of T1 heterozygous plants was used to pollinate self-stigmas, *At-PrsS*_*1*_ or Col-0 stigmas. Pollen transmission was examined by checking the GFP signals (resulting from the pFAST-Green vector backbone) in the seeds. The ratios of GFP positive seeds of each pollination are shown in the table. The actual numbers of GFP positive seeds and total seeds in each pollination are indicated in the brackets. For T2 seeds of heterozygous *At-SI/hld1/cHLD1* lines, a ∼50% of GFP positive ratio was observed, whereas a ratio of ∼75% was observed for *At-SI/hld1/cHLD1(S218A)* lines **(Fig 3C; Table S4)**. We reasoned that for the heterozygous T1 *At-SI/hld1/cHLD1* lines, expression of *cHLD1* in the *hld1* mutant pollen rescued the *hld1* mutant phenotype, resulting in SI-induced pollen death, and thus prevented pollen containing the *cHLD1* transgene from achieving fertilization, which led to the reduction of GFP positive T2 seeds ratio from theoretical 75% to ∼50%. However, for the heterozygous T1 *At-SI/hld1/cHLD1(S218A)* lines, the S218A mutation could not rescue the *hld1* mutant deficiency, therefore resulting in normal Mendelian segregation of GFP signals in the T2 seeds. Consistent with this hypothesis, there is a significant reduction in the GFP positive seeds ratio when *At-SI/hld1/cHLD1* pollen was pollinated onto *At-PrsS*_*1*_ stigmas compared with that of ♀*Col-0* x ♂*At-SI/hld1/cHLD1* **(Fig 3C; Table S4)**. However, this difference of GFP positive seeds ratio was not observed for *At-SI/hld1/cHLD1(S218A)* lines **(Fig 3C; Table S4)**. These data showed that the *HLD1* WT allele rescued the *hld1* mutant phenotype, but not the S218A mutant allele.

| Line | Selfing | ♀ *At-PrsS<sub>1</sub>* | ♀ *Col-0* |
| --- | --- | --- | --- |
| *At-Sl/hld1-7/pHLD1::mCherry-cHLD1*<sup>+/-</sup> line 4 | 46.8%(109/233) | - | - |
| *At-Sl/hld1-7/pHLD1::mCherry-cHLD1*<sup>+/-</sup> line 11 | 48.1%(115/239) | 4.7%(25/532) | 67.1%(457/681) |
| *At-Sl/hld1-7/pHLD1::mCherry-cHLD1*<sup>+/-</sup> line 16 | 48.5%(113/233) | 2.6%(16/604) | 73.2%(510/697) |
| *At-Sl/hld1-7/pHLD1::mCherry-cHLD1*<sup>+/-</sup> line 19 | 50.2%(139/277) | 1.9%(8/432) | 74.0%(273/369) |
| *At-Sl/hld1-7/pHLD1::mCherry-cHLD1*<sup>+/-</sup> line 21 | 56.5%(210/372) | - | - |
| *At-Sl/hld1-7/pHLD1::mCherry-cHLD1*<sup>+/-</sup> line 24 | 52.9%(156/295) | - | - |
| *At-Sl/hld1-7/pHLD1::mCherry-cHLD1(S218A)*<sup>+/-</sup> line 1 | 75.6%(198/262) | - | - |
| *At-Sl/hld1-7/pHLD1::mCherry-cHLD1(S218A)*<sup>+/-</sup> line 2 | 75.3%(226/300) | - | - |
| *At-Sl/hld1-7/pHLD1::mCherry-cHLD1(S218A)*<sup>+/-</sup> line 6 | 72.2%(156/216) | 49.1%(112/228) | 50.1%(305/602) |
| *At-Sl/hld1-7/pHLD1::mCherry-cHLD1(S218A)*<sup>+/-</sup> line 8 | 77.1%(269/349) | - | - |
| *At-Sl/hld1-7/pHLD1::mCherry-cHLD1(S218A)*<sup>+/-</sup> line 12 | 76.1%(249/327) | - | - |
| *At-Sl/hld1-7/pHLD1::mCherry-cHLD1(S218A)*<sup>+/-</sup> line 16 | 76.0%(187/246) | 58.2%(102/175) | 49.5%(103/208) |
| *At-Sl/hld1-7/pHLD1::mCherry-cHLD1(S218A)*<sup>+/-</sup> line 23 | 80.5%(186/231) | 52.4%(204/389) | 51.1%(218/427) |

**Supplemental Table S5.**
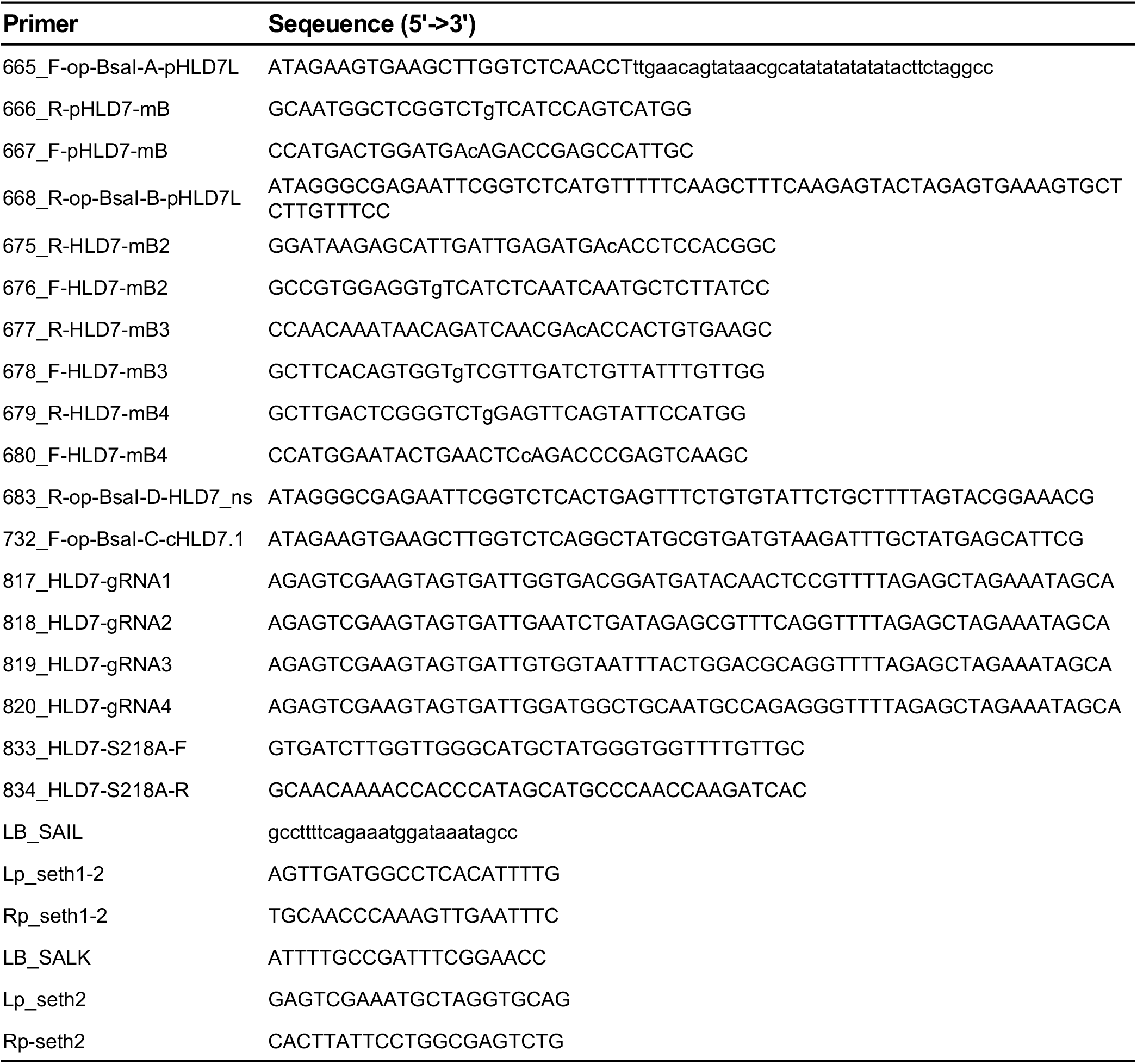
Primers used in this research.

